# Integrative genomics approach identifies glial transcriptomic dysregulation and risk in the cortex of individuals with Alcohol Use Disorder

**DOI:** 10.1101/2024.08.16.607185

**Authors:** Anna S. Warden, Nihal A. Salem, Eric Brenner, Greg T. Sutherland, Julia Stevens, Manav Kapoor, Alison M. Goate, R. Dayne Mayfield

## Abstract

Alcohol use disorder (AUD) is a prevalent neuropsychiatric disorder that is a major global health concern, affecting millions of people worldwide. Past molecular studies of AUD used underpowered single cell analysis or bulk homogenates of postmortem brain tissue, which obscures gene expression changes in specific cell types. Here we performed single nuclei RNA-sequencing analysis of 73 post-mortem samples from individuals with AUD (N=36, N_nuclei_= 248,873) and neurotypical controls (N=37, N_nuclei_= 210,573) in both sexes across two institutional sites. We identified 32 clusters and found widespread cell type-specific transcriptomic changes across the cortex in AUD, particularly affecting glia. We found the greatest dysregulation in novel microglial and astrocytic subtypes that accounted for the majority of differential gene expression and co-expression modules linked to AUD. Analysis for cell type-specific enrichment of aggregate genetic risk for AUD identified subtypes of microglia and astrocytes as potential key players not only affected by but causally linked to the progression of AUD. These results highlight the importance of cell-type specific molecular changes in AUD and offer opportunities to identify novel targets for treatment.

## Introduction

Alcohol Use Disorder (AUD) is a prevalent neuropsychiatric disorder characterized by impaired ability to stop or control alcohol use despite adverse physical, psychological, or sociological consequences. Worldwide over 100 million people had an alcohol use disorder in the last year and repeated alcohol abuse constitutes the seventh leading risk factor for death and disability worldwide(1,2). Specifically, in the United States, 28.6 million adults ages 18 and older (11.3% of age group) had AUD in the past year(3). Despite its prevalence there are currently only a handful of treatments for treating AUD, with minimal to moderate efficacy in reducing harmful alcohol consumption. To develop more effective therapeutic targets, we must elucidate how alcohol combines with genetic risk to converge onto specific molecular processes that lead to the development and progression of AUD.

Although AUD is highly heritable, its genetic etiology is complex with additive effects of alcohol, combined with behavioral, environmental, psychological, and physiological factors(4). Assessment of AUD risk is also challenging due to its genetic architecture, complexity of alcohol behaviors and the dynamic patterns of alcohol drinking behavior across lifespan(4). However, despite this genetic complexity, molecular studies have identified consistent patterns of gene expression changes in post-mortem brain tissue from AUD individuals(5,6). At the transcriptomic level, AUD brains exhibit region-dependent alterations of genes involved in neuronal activity with a concomitant upregulation of genes involved in inflammation(5–8). Of note, most of these transcriptomic studies in human brain have been bulk analyses of tissue homogenates, which may obfuscate or minimize cell-type specific gene expression patterns, particularly in minor cell populations.

Developments in sequencing technology, such as single nucleus RNA sequencing (snRNAseq), have permitted unprecedented resolution into the molecular architecture of the human brain(9,10). Previously, we profiled a small cohort of postmortem human dorsolateral prefrontal cortex (dlPFC) samples using snRNAseq(11). Despite the small sample size, this study identified novel differentially expressed genes and found an enrichment of inflammatory genes in multiple glial subtypes in a region that is key in reward processing and the pathophysiology of addiction(11). Similarly, a recent snRNAseq study in the nucleus accumbens of individuals with AUD, identified neuroinflammatory changes in glia that were underrepresented in previous bulk transcriptomes. Limitations of these studies include the number of nuclei sequenced, size of the cohorts (<10 per group) and representation of a single sex.

In this study we leveraged snRNAseq from postmortem human dlPFC of a cohort of 36 individuals with AUD and 37 control individuals for a total of 73 samples across two institutions to define the transcriptomic architecture of AUD across 32 cell type classes. We identified novel differentially expressed genes between cell populations in AUD, determined co-expression modules that correlated with DSM-5 diagnosis and identified key hub genes that may be implicated in the pathogenesis of AUD. The largest number of differentially expressed genes were detected in astrocytes, oligodendrocytes, and microglia. Finally, by integrating GWAS summary statistics we highlight causal cell types, with most of the enrichment found in glial subtypes. This study provides the first comprehensive, well-powered study of the broad transcriptional changes in human brain in AUD on the single nuclei level and further highlights the clinical relevance of glial and immune changes in both the pathogenesis and progression of AUD risk.

## Materials & Methods

Additional information is available in supplemental methods.

### Cohort selection and postmortem tissue collection

Diagnosis of alcohol dependence was denoted as based on DSM-5 criteria. Postmortem brain samples were provided by the New South Wales Brain Tissue Resource Centre at the University of Sydney. Cohort selection and processing of fresh frozen tissue samples were collected as previously described(8) (Table S1).

### Droplet-based single-nucleus RNA-seq

For droplet-based single-nucleus RNA sequencing (snRNA-seq), libraries were prepared as previously described[8]. Files were processed using Cellranger (v6.1.2) and aligned to the GRCh38 reference with default parameters before downstream processing.

### Filtering, normalization & visualization

Ambient RNA was corrected using SoupX(v1.3.0) as previously described(12). The corrected matrix was then loaded into Seurat(v4.2.0). Custom filtering thresholds and normalization can be found in supplemental methods. Due to batch effects between sequencing sites, we next integrated our cohort using Harmony(v1.1.0)(13) and then followed a standard Seurat pipeline. Normalized expression of every gene was centered to a mean of zero. Principal component (PC) analysis was performed on highly variable genes, the top 20 PCs were used to construct a nearest neighbor graph. Nuclei were then assigned to clusters using the Louvain algorithm with the resolution set to 0.5.

### Marker identification for cell type specific clusters

We characterized clusters for a set of marker genes defined by differential expression analysis of the cells group in each cluster against the remaining cells within the corresponding aggregate cell type cluster using Seurat FindMarkers function. This analysis was applied to all clusters independently. Significantly over-expressed genes were defined based on the Wilcoxon-rank-sum test with an FDR<0.05, |log2FC|>0.5. Only genes detected in at least 25% of the cells within the given subcluster were considered. Gene ontology (GO) enrichment analysis was performed using Metascape(14). Detailed cluster annotation can be found in supplemental methods.

### Cell type proportion analysis

To determine if AUD changed proportions of cell type-specific nuclei we used the speckle package (v 3.18) propeller method in R environment(15) as well as scProportionTest (v0.0.0.9000) in R environment (https://github.com/rpolicastro/scProportionTest). Results were filtered based on log2FC>|0.2|, FDR<0.05.

### Differential gene expression analysis

DE testing was performed using the libra pseudobulking package(v1.0.0)(16). First, nuclei corresponding to a given aggregate cell type were selected from the full dataset. Second, counts were summed to produce a pseudobulk matrix for each cell type. Lastly, DEGs between alcohol-dependent and control individuals were detected using edgeR(v3.36.0) with correction for the following covariates: Sex, Age, PMI, AUD Classification and Sequencing Site. FDR<0.05 was used to detect DEGs within each cell type. Consistency was determined using a resampling approach, as previously described(17).

### High dimensional weighted gene correlation network analysis

Network analysis was performed with hdWGNCA(v0.1.1), a comprehensive framework for analyzing co-expression networks in high dimensional transcriptomics data such as single-cell and spatial RNA-seq(18,19). A soft power of 7 and a minimum module size of 25 genes was selected. This analysis included gene network construction, module-trait correlation, module eigengene by trait, and hub gene analysis. We selected DSM-5 categorical variables and identified eigengene expression of each module for these variables in addition to AUD Classification.

### External data sources

Data sources for glial subtypes can be found in Table S9 (see supplemental methods for details) (20–27) . Human AUD dlPFC datasets: bulk RNAseq(8); proteome(28)

### GWAS enrichment meta-analysis

Summary statistics for meta-analysis of alcohol dependence and AUDIT-C GWAS were obtained from the Million Veteran Program (MVP) (N=318,694)(29). MAGMA(v1.10)(30,31) was used to perform gene-based association analysis for the GWAS summary statistics as previously described(8,11). We selected the European ancestry summary statistics since 72 out of 73 individuals in our study are of European descent. The output of gene-based analysis was used to perform the cell-type enrichment analysis for the GWAS genes. The enrichment scores were calculated on (1) cluster specific DEGs in AUD individuals (FDR<0.05); (2) aggregate DEGs in major cell type classes in AUD individuals (FDR<0.05); (3) the ranked genes by kME of each hdWGCNA module.

### Statistical analysis & Graphing

Module eigengene statistical comparisons were performed by two-tailed Welch’s t-test (P<0.05). Significance of overlapping gene sets was performed using a one-sided hypergeometric test on gene sets (P<0.05). Graphs for figures were produced using R(v4.3.1) or Prism(v10.1.0) (GraphPad, San Diego, CA).

### Ethical approval

The Authors are in support of general ethics and inclusion guidelines for researchers in multi-region collaborations involving local researchers to promote greater equity in research.

### Data & Code Availability

Original code and scripts are available upon request.

## Funding

National Institutes of Health: U01-AA020926 to R.D.M., R01-AA012404 to R.D.M. Collaborative Studies on Genetics of Alcoholism: 5 U10 AA08401 to A.M.G.

### Contributions

R.D.M. and A.M.G. planned the study. G.T.S. and J.S. provided the postmortem brain samples from the NSW Tissue Resource Centre. E.B., M.K, and N.A.S. performed the isolations and acquired the data for analysis. N.A.S. and E.B. performed the initial QC analysis and integration. A.S.W. performed the data analysis and wrote the manuscript. R.D.M and N.A.S. provided data review and key insights for subsequent analyses. R.D.M, N.A.S, G.T.S., J.S., A.M.G. and M.K. provided critical comments and suggestions on the manuscript.

## Results

To investigate AUD-specific transcriptional disruption within the complex architecture of the human brain, we used snRNAseq to profile postmortem tissue samples from the dlPFC (Fig 1A). We curated a cohort of 36 individuals with AUD and 37 control individuals for a total of 73 samples across two institutions including both sexes: (9 AUD females, 10 control females; 27 AUD males, 27 control males), a range of ages (24-81 years, average 54.5 years) and postmortem interval (8.5-72 hours, average 30.8 hours) (Fig 1A). Because sequencing occurred between two institutional sites, in depth quality control analyses were performed to ensure that neither set of samples were skewed for any clinical metadata covariates or snRNAseq quality metrics before pooling (Fig S1). Complete clinical and sequencing metadata can be found in Table S1. We corrected for ambient RNA contamination using SoupX(12) and removed mitochondrial contamination above a set threshold using Seurat, resulting in 459,446 nuclei. We visualized the data using uniform manifold approximation and projection (UMAP) and integrated datasets across sites using Harmony(13) (Fig S2).

**Figure 1:**
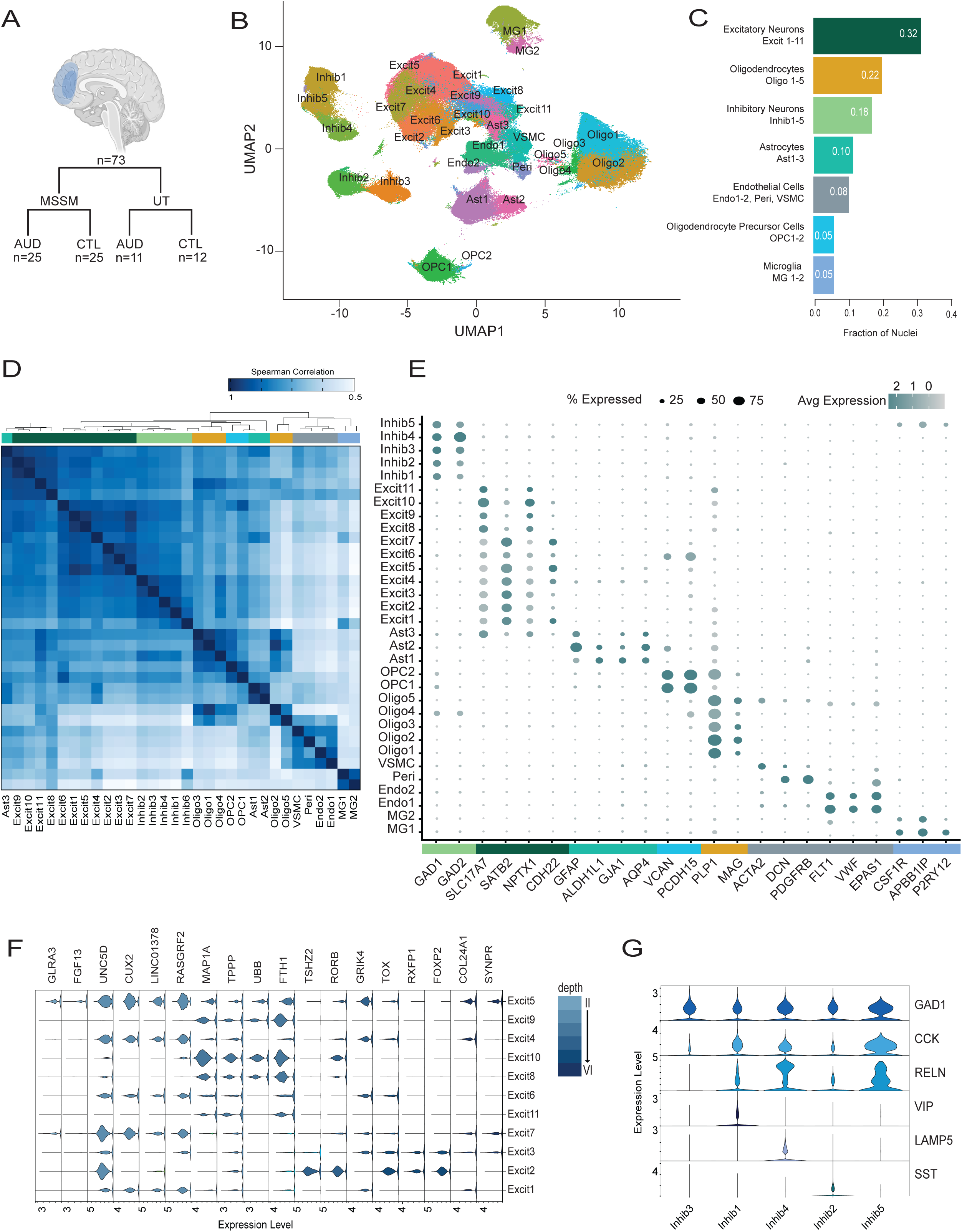
Cell type identification in the dlPFC of Control and AUD individuals. (A) Schematic of sample cases and sequencing sites for the nuclei isolated from 73 brains. (B) UMAP plots of the 459,446 nuclei in our dataset demonstrating transcriptomically distinct cell type clusters determined by unsupervised clustering. (C) Fraction of nuclei represented by specific cell types/clusters. (D) Spearman correlation heatmap between clusters demonstrates transcriptomically distinct cell types and clustering. The color intensity represents the correlation value between 0.5 and 1. (E) Scaled mean expression levels of known cell type markers in the 32 clusters shown as a stacked dot plot. The size of the dots represents the proportion of cells expressing the gene (Pct. exp.), whereas the color intensity represents the average expression level (Avg. exp. scale) (F) Cortical-layer-specific markers varied in expression within the excitatory neuronal clusters. The violin plots depict the expression per cluster of layer-specific marker genes, from the more superficial layers (II) on the left to the deeper layers (V/VI) on the right. The color intensity represents the layer depth. (G) Known classes of inhibitory neurons were identifiable based on the expression pattern of inhibitory marker (GAD1), peptide genes (VIP, SST and CCK) and interneuron markers (RELN, LAMP5).

### Identification of refined subpopulations in human alcohol-dependent dlPFC

Transcriptionally distinct groups of nuclei were identified using unsupervised, graph-based clustering which yielded 32 clusters in the combined population of AUD and control subjects (Fig 1B). The 32 clusters captured all major cell types of the dlPFC: subtypes of excitatory and inhibitory neurons, microglia, astrocytes, oligodendrocyte precursor cells (OPCs), oligodendrocytes, endothelial cells, pericytes and vascular smooth muscle cells (VSMCs) (Fig 1B). Proportions of different cell types were comparable to other snRNA-seq studies(11,17) and similar across samples and sex, with neurons being over-represented relative to glia (50% neurons, 22% oligodendrocytes, 10% astrocytes, 8% endothelial cells, 5% microglia, 5% OPCs) (Fig 1C, Fig S3). Spearman correlation of average expression between cell type-specific clusters showed tight correlation of neuronal populations and distinct separation of non-neuronal cell types (Fig 1D). We verified these annotations based on known marker gene expression patterns from single cell/nuclei studies from the prefrontal cortex(17,32–36) (Fig 1E). Gene expression patterns previously linked to specific cortical layers matched our clustering of excitatory neurons, with cluster specific representation across layers I-VI (Fig 1F) (32). Likewise, inhibitory cell types demonstrated cluster specific interneuron gene expression patterns similar to other snRNAseq studies in human dlPFC in other neuropsychiatric disorders (Fig 1G) (32) .

### Defining glial subpopulations in AUD

We hypothesized that AUD leads to relative changes in the abundance of specific cell types as a consequence of disease pathology, similar to other neurological diseases(15,37). We observed changes in proportions of specific clusters between AUD and controls specifically in microglia (MG1-2), astrocytes (Ast1-3), and oligodendrocytes (Oligo 2,3,5) (Fig 2, Table S3). There were shifts in proportions between two microglial subclusters, with a decrease in MG1 (log2FC: -0.40, FDR=0.00099) and an increase in MG2 (log2FC: 2.41, FDR=0.00099) in AUD (Fig 2A-B, Table S3). To define the identity of those subclusters, we utilized known microglia state markers and performed gene ontology enrichment(20–22). MG1 was enriched in known homeostatic markers and functions (WP304, -log_10_ P=11.11) (Fig 2C-D, Table S4). MG2 in contrast, was enriched in disease associated microglia (DAM) (P<1.06e^-63^) as well as border associated macrophage (BAM) markers (P < 0.014) (20–22). Additionally, MG2 was enriched in pathways related to immune signaling (GO:0071345, -log_10_ P=17.44) (Fig 2C,E, Table S4). These inflammatory and DAM/BAM markers suggest a shift to an diseased microglial phenotype in AUD, consistent with but expanding on previous studies highlighting the role of inflammatory signaling in microglia in regulation of AUD phenotypes(38–40).

**Figure 2:**
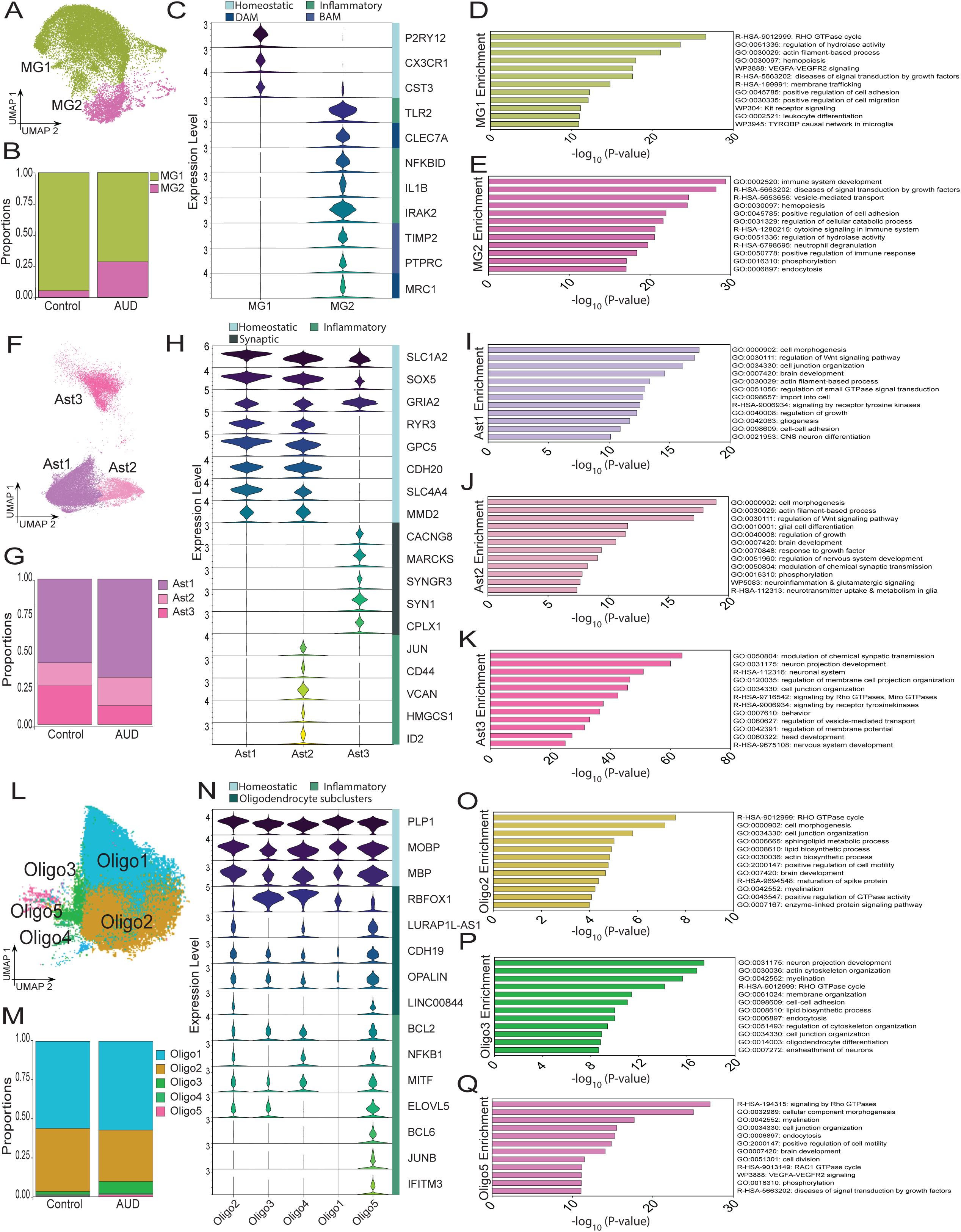
Cell type specific clustering reveals alcohol-responsive glial clusters in AUD. (A) UMAP plot of microglia clusters 1 (MG1) and 2 (MG2). (B) Microglia cluster proportions between Control (MG1: 0.94, MG2: 0.05) and AUD (MG1: 0.71, MG2: 0.28). (C) Violin plot of microglia subtype markers in MG1 and MG2: homeostatic markers (P2RY12, CX3CR1, CST3), inflammatory markers (TLR2, NFKBID, IL1B, IRAK2), disease-associated microglia markers (CLEC7A, MRC1), and border associated macrophage markers (PTPRC, TIMP2). Gene ontology enrichment for alcohol-responsive clusters (D) MG1 and (E) MG2 for the top 12 categories. Enrichment is a function of -log10 (P-value). (F) UMAP plot of astrocyte clusters 1 (Ast1), 2 (Ast2), and 3 (Ast3). (G) Astrocyte cluster proportions between Control (Ast1: 0.58, Ast2: 0.15, Ast3: 0.27) and AUD (Ast1: 0.67, Ast2: 0.20, Ast3: 0.13). (H) Violin plot of astrocyte subtype markers in Ast1-3: canoncial markers (SLC1A2, SOX5, GRIA2), Ast1-2 specific astrocyte markers (RYR3, GPC5, CDH20, SLC4A4, MMD2), synaptic-astrocytic markers specific to Ast3 (CACNG8, MARCKS, SYNGR3, CPLX1), and inflammatory markers specific to Ast2 (JUN, CD44, VCAN, HMGCS1, ID2). Gene ontology enrichment for alcohol-responsive clusters (I) Ast1, (J) Ast2 and (K) Ast3 for the top 12 categories. Enrichment is a function of -log10 (P-value). (L) UMAP plot of oligodendrocyte clusters 1-5 (Oligo1-5). (M) Oligodendrocyte cluster proportions between Control (Oligo1: 0.57, Oligo2: 0.40, Oligo3: 0.017, Oligo4: 0.009, Oligo5: 0.004) and AUD (Oligo1: 0.57, Oligo2: 0.33, Oligo3: 0.08, Oligo4: 0.01, Oligo5: 0.01). (N) Violin plot of oligodendrocyte markers in Oligo1-5: myleination markers (PLP1, MOBP, MBP), subtype markers (RBFOX1, LURAP1L-AS1, CDH19, OPALIN, LINC00844), and inflammatory/disease-associated markers (BCL2, NFKB1, MITF, EVOLV5, BCL6, JUNB, IFITM3). Gene ontology enrichment for alcohol-responsive cluster (O) Oligo2, (P) Oligo3, and (Q) Oligo5 for the top 12 categories. Enrichment is a function of -log10 (P-value). For the violin plots in C, H, N values extend from minimum to maximum, color is a function of individual genes.

There were also shifts in astrocyte populations in AUD (Ast1 log2FC=0.23, FDR=0.00099; Ast2 log2FC=-0.36, FDR=0.00099; Ast3 log2FC=-1.08, FDR=0.0099) (Fig 2F-G, TableS3). We first characterized these clusters according to known astrocyte subpopulation markers(41,42). Ast1 was *SLC1A2*^+^/SOX5^+^ and enriched in canonical astrocyte functions (GO:0042063, -log_10_ P=11.71) (Fig 2H-I, FigS4, Table S4). Ast2 also expressed canonical markers (*SLC1A2^+^/SOX5*^+^) but was enriched in neuroinflammatory pathways (WP5083, -log_10_ P=7.68) (Fig 2H,J, Fig S4, Table S4) (42). This is consistent with the role of astrocytes in regulation of AUD(40,43,44) as well as previous snRNAseq studies in AUD(11,45). Ast3, in addition to expressing canonical astrocytic markers, also expressed unique and novel synaptic markers specific to the recently reported synaptic neuron and astrocyte program (SNAP-a) (Fig 2H, Fig S4A)(46). This gene expression program involves concerted effects on the expression of distinct genes for synaptic components in neurons and astrocytes as well as homeostatic astrocytic functions(46). Ast3 GO category enrichment was equally split between canonical astrocyte- and synaptic-specific functions such as modulation of chemical synaptic transmission (GO:0050804, -log_10_ P=67.30) (Fig 2K, Fig S4F, TableS4). SNAP-a GO category enrichment also showed strong overlap with Ast3 (Fig S4G). Expression of SNAP-a genes with our astrocyte clusters also showed specific overlay in Ast3, suggesting regulated neuron-astrocyte programs in our astrocytic subclusters (Fig S4H-I). The expression of SNAP-a programs in AUD has not previously been reported and the decreased expression of SNAP-a in AUD suggests a novel loss of homeostatic control of synaptic regulation in favor of inflammatory functions in astrocytes that may play a key role in AUD pathogenesis.

Proportions of oligodendrocyte subpopulations also shifted between AUD and controls individuals (Oligo2 log2FC= -0.29, FDR=0.0016; Oligo3 log2FC=2.117, FDR=0.0016; Oligo5 log2FC=1.59, FDR=0.00099) (Fig 2L-M, Table S3). All oligodendrocyte clusters had strong enrichment of myelination genes (i.e. *PLP1, MOBP, MBP*) in both AUD and control individuals (Fig 2N) (GO:0042552, Oligo2, -log_10_ P=19.73; Oligo3, -log_10_ P=15.97; Oligo5, -log_10_ P=17.71) (Table S4). Oligo5 was enriched in immune-related genes such as *NFKB1* and *IFITM3* (P<0.031) and was proportionally increased in AUD (Fig 2M,Q). These markers are novel in AUD and are additionally consistent with recent work demonstrating the inflammatory subpopulations of oligodendrocytes in the white matter of individuals with multiple sclerosis(47,48). Additionally, subclusters were separately enriched in different oligodendrocyte functions such as lipid biosynthesis (GO:0008610, Oligo2, -log_10_P=8.81; Oligo3, -log_10_ P=9.10), and diseases of signal transduction (R-HSA-5663202, Oligo5, -log_10_ P=11.089) (Fig 2O-Q, Table S4). Taken together, these AUD-responsive changes in oligodendrocytes, microglia and astrocytes, suggest glial populations may drive AUD pathogenesis.

### Glial clusters are primary drivers of the human AUD transcriptional signature

To identify differentially expressed genes (DEGs) we performed pseudobulking, including covariates to account for age, sex, PMI, AUD classification and sequencing site, as previously described(16,49). We identified 3667 DEGs across all cell types (Padj<0.05), with 1875 upregulated and 1792 downregulated in AUD (Fig 3A, Table S5). The majority of DEGs were identified in glial populations (Fig 3A). We additionally tested consistency between our single nuclei DEGS and bulk expression, finding strong consistency at a cell type-specific level to previous findings (Fig 3B) (8).

**Figure 3:**
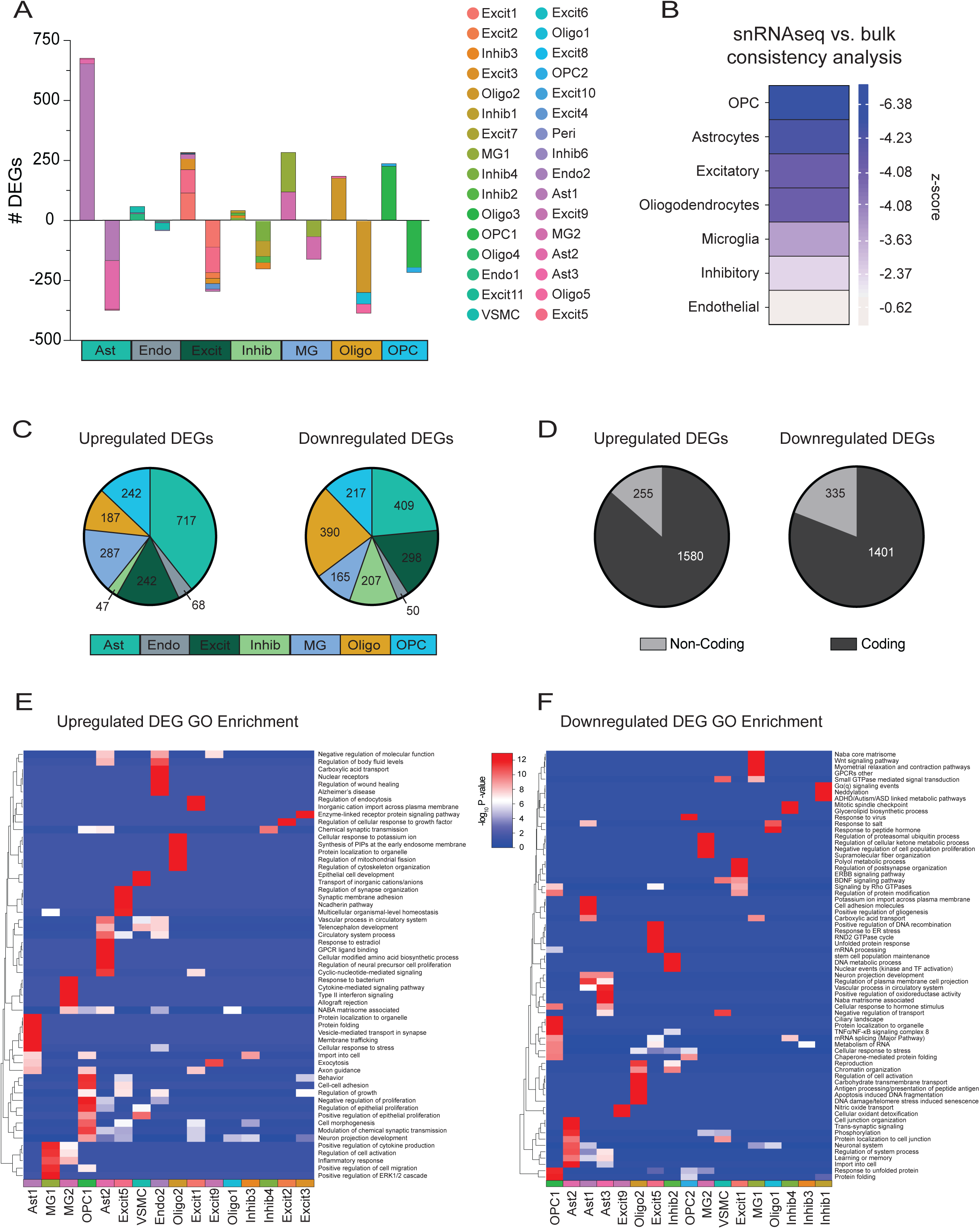
Glial genes drive differential expression in AUD. (A) Number of differentially expressed genes (DEGs) per cluster, with a total of 3667 DEGs. The y-axis represents the number of DEGs, positive values indicate upregulated genes and negative values represent downregulated genes. Colors represent specific clusters which map to the legend on the right. Aggregate cell type classifications are mapped on the x-axis. (B) Global consistency analysis of snRNA-seq versus bulk RNA-seq. Agreement or disagreement was estimated by deviation from random expectation (*z*-score) of single-cell DEG average rank scores in the ranked list of genes from bulk differential analysis. (C) Circle chart for upregulated and downregulated DEGs. Aggregate cell type classes are represented by color and map to the legend on the bottom of the graphs. Numbers represent the number of DEGs per aggregate cell type cluster. (D) Circle charts for upregulated and downregulated DEGs as a function of protein-coding versus non-coding. GO enrichment of (E) upregulated DEGs or (F) downregulated per cell type represented as a heatmap. Color represents enrichment as a function of -log_10_ P-value, with red indicating strong enrichment, light blue representing weak enrichment and dark blue representing no enrichment. Specific cell type clusters are labeled on the X-axis and map to the UMAP legend.

Specific glial clusters represented the majority of upregulated DEGs, such as Ast1, MG1 and Oligo2 (Fig 3C). Excitatory neurons had the second largest fraction of upregulated DEGs after astrocytes, similar to our pilot study (Fig 3C)(11). Interestingly, approximately 17% of upregulated DEGs were non-coding genes (Fig 3D), reinforcing their known role in AUD(5,50). Gene ontology and pathway enrichment of upregulated DEGs per cell type reinforced the role of inflammatory signaling in AUD in glial cell types, especially microglia (Fig 3E, Table S6). Microglia-specific DEGs were enriched in canonical immune response (GO:0001819, -log_10_P=8.04) in addition to novel viral response pathways including type II interferons (WP619, -log_10_P=5.59). Astrocyte-specific upregulated DEGs showed strong enrichment in regulation of cellular response to stress (R-HSA-2262752, -log_10_P=7.25) and the extracellular matrix (ECM) (R-HSA-1474244, -log_10_P=3.71) (Fig 3E, Table S6). Excitatory and inhibitory neuron upregulated DEGs had strong enrichment in regulation of the synapse (GO:0050807, -log_10_P=4.63; GO:0007268, -log_10_P=3.18) (Fig 3E, Table S5). Moreover, the majority of DEGs and pathway enrichments were unique to each cell type, suggesting that key processes such as inflammation overlap between glial cell types in AUD but transcriptional shifts in AUD are primarily cell type-specific (Fig 3E, Table S5-6). Many, but not most, of these patterns of DEGs have been previously implicated in AUD and in regulation of alcohol behaviors(51).

Downregulated DEGs were distributed evenly across cell type specific clusters (Fig 3A,C, Table S5). Glial clusters represented the majority of downregulated DEGs with excitatory neurons having the third largest fraction after astrocytes and oligodendrocytes (Fig 3A,C, Table S5). Approximately 23% of downregulated DEGs represented non-coding genes (Fig 3C). Astrocyte downregulated DEGs were enriched in gliogenesis (GO:0042063, -log_10_P=4.13) and trans-synaptic signaling (GO:0099537, -log_10_P=6.27), suggesting dysregulation of astrocyte populations and synaptic regulatory function in AUD (Fig 3F, Table S6). Microglial downregulated genes were enriched in regulation of cell proliferation (GO:0008285, -log_10_P=3.48) and lipid biosynthetic process (GO:0046467, -log_10_P=2.12) (Fig 3F, Table S6). Oligodendrocyte downregulated genes were enriched in senescence (R-HSA-2559586, -log_10_P=9.32) (Fig 3F, Table S6). Excitatory neurons and inhibitory neurons downregulated genes were enriched in chromosome maintenance and organization (R-HSA-73886, -log_10_P=2.07; GO:0006325, -log_10_P=3.22). Downregulated DEGs had some cell-type specific patterns but had more overlap between core homeostatic functions downregulated in AUD such as cellular responses to stress and protein folding (Fig 3F, Table S6).

### Microglia high dimensional WGCNA modules identify inflammatory and immune targets in AUD

Because the majority of differentially expressed genes in AUD were in glia, we chose to further identify disease and cell type-specific gene co-expression network changes in AUD in selected cell types. We leveraged high dimensional weighted gene correlation network analysis (hdWGCNA) for each aggregate cell type. hdWGCNA is a framework for co-expression network analysis in single-cell/nuclei and spatial transcriptomics data designed to identify disease-relevant co-expression network modules in specific cell populations(18,19).

Microglia clusters MG1 and MG2 were combined for hdWGCNA analysis in order to identify both cluster specific and microglial changes. The microglial network contained 8 modules, 3 of which had increased eigengene expression in AUD (Fig 4A, Table S7). Microglia module MG-M2, 360 genes, associated with cluster MG1, was more highly expressed in mild and moderate AUD (Fig 4B-E). Module MG-M2 was also weakly positively correlated with total drinking years (R=0.19) (Fig 4C, Table S8). Module MG-M2 was highly enriched in DAM (-log_10_P=10.07) and interferon response microglia (IRM) (-log_10_P=6.95) (Fig 4F, Table S9)(21,22). In concordance with the microglia subtype enrichment module MG-M2 was enriched in canonical immune functions such as cytokine signaling (GO:0071345, -log_10_P=14.80) and lipid production (hsa05417, -log_10_P=10.07) in addition to novel pathways like interferon signaling (GO:0032729, -log_10_P=4.51) (Fig 4G, Table S10). Top hub genes for module MG-M2 included known immune targets for AUD such as *TLR2* and *IL1B* as well as other components of NFkB signaling such as *NFKBIA* and *IRAK2* (Fig 4H)(5,51,52).

**Figure 4:**
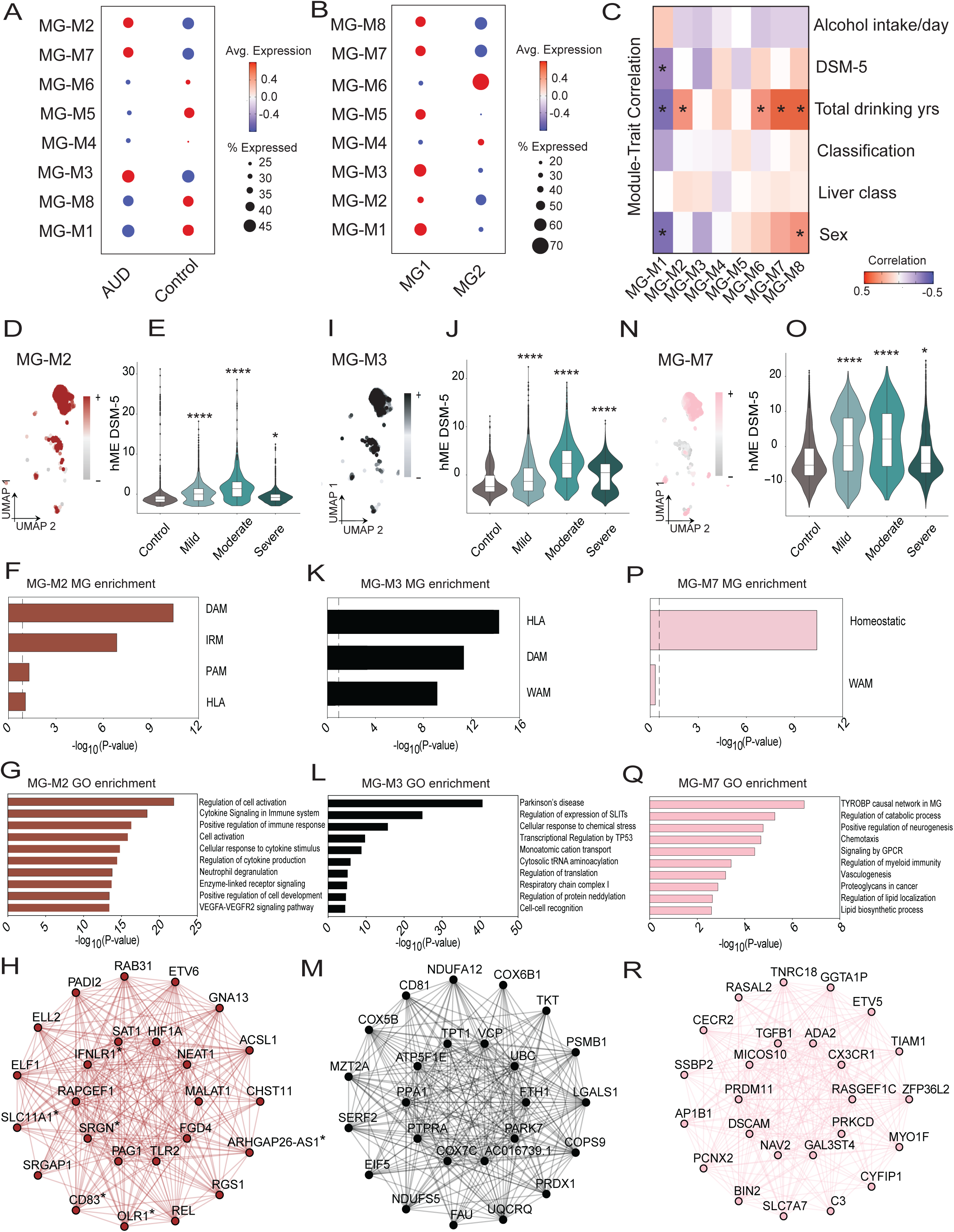
Microglia co-expression modules highlight immune dysfunction and loss of homeostatic regulation in AUD. Co-expression modules from microglia clusters represented by dot plot as the module eigengene (hME) average expression (color of dot) and the percent of genes expressed in each module (size of dot) as a function of (A) AUD classification or (B) cell type cluster. Red indicates increased average expression, blue indicates decreased average expression of the module. (C) Module-trait correlation heatmap for specific traits: Alcohol intake (g/day), DSM-5, Total drinking years, AUD Classification, Liver class and Sex. Color is a function of correlation strength, with red representing positive correlation and blue representing negative correlation with module eigengene expression. *R ≥ |0.2|. (D) UMAP plot colored by modules eigengenes for microglia co-expression module MG-M2. (E) hME expression of microglia co-expression module MG-M2 across DSM-5 categories. (F) MG-M2 enrichment for microglia subtypes (DAM, IRM, PAM and HLA). (G) Top 10 GO category enrichments for MG-M2. (H) Visualization of top hub genes for MG-M2. Genes marked with an asterisk (*) represent hub genes that are also differentially expressed in microglia clusters. (I) UMAP plot colored by modules eigengenes for MG-M3. (J) hME expression of microglia co-expression module MG-M3 across DSM-5 categories. (K) MG-M3 enrichment for microglia subtypes (HLA, DAM, WAM). (L) Top 10 GO category enrichments for MG-M3. (M) Visualization of top hub genes for MG-M3. (N) UMAP plot colored by modules eigengenes for microglia co-expression module MG-M7. (O) hME expression of microglia co-expression module MG-M7 across DSM-5 categories. (P) MG-M7 enrichment for homeostatic microglia subtype. (Q) Top 10 GO category enrichments for MG-M7. (H) Visualization of top hub genes for MG-M7. Enrichment is a function of -log10 (P-value). For the violin plots in E, J, O values extend from minimum to maximum, with a white boxplot to show the median value. Color is a function of DSM-5 category. Two-tailed Welch’s t-test for DSM-5 module eigengene expression comparisons, * P<0.05, **P<0.01, ***P<0.001, ****P<0.0001.

Similarly, module MG-M3, 103 genes, was highly expressed in cluster MG1 and significantly increased in mild, moderate and severe AUD (Fig 4A,B,I,J). Module MG-M3 overlapped with antigen-presenting response microglia (HLA) (-log_10_P=13.49), DAM (-log_10_P=11.64) and white matter-associated microglia (WAM) (-log_10_P=8.82) (Fig 4K, Table S9)(21,22,24). Module MG-M3 also had strong enrichment in Parkinson’s disease (hsa05012, -log_10_ P =40.66) and cellular response to stress (R-HSA-9711123, -log_10_P=15.75) (Fig 4L, Table S9). Hub genes included neurodegenerative targets such as *PARK7*, which is a potential biomarker of AUD in humans (Fig 4M, Table S7)(53).

Module MG-M7, 79 genes, was expressed in cluster MG1 and upregulated in mild and moderate AUD (Fig 4A,B,N,O, Table S7). Module MG-M7 positively correlated with total drinking years (R=0.25) (Fig 4C, Table S8). Module MG-M7 only overlapped with homeostatic microglia markers (-log_10_P=10.04) (Fig 4P, Table S9)(54). Moreover, module MG-M7 was enriched in canonical microglia functions such as TYROBP causal network (WP3945, -log_10_P=6.51) and complement system (WP5090, -log_10_P=2.70) (Fig 4Q, Table S10). Complement signaling is highly implicated in synapse elimination and pruning in the brain, suggesting a regulatory role of microglia in synapse elimination in AUD(55,56). Hub genes included common homeostatic and complement markers (*TGFB1*, *CX3CR1*, *C3*) (Fig 4K, Table S7). The identification of microglia subtypes in AUD, common to other neurological disorders such as DAM populations, suggests a novel core disease state of microglia in AUD(57). Furthermore, identified co-expression pathway targets such as interferon signaling in microglia are promising since interferon signaling regulates many of the previously implicated neuroimmune targets in AUD that regulate behavior including TLR3 and TLR7 signaling (58–60).

### Astrocyte and oligodendrocyte modules highlight roles for synapse regulation and cellular stress in AUD

We additionally constructed networks for the other major glial cell types, the astrocyte network consisted of four modules, all of which had increased eigengene expression in AUD (Fig 5A, Table S7). Of interest was module Ast-M1, containing 2005 genes, primarily expressed in the SNAP-a cluster, Ast3 (Fig 5B,D). Module Ast-M1 was upregulated in mild, moderate and severe AUD (Fig 5E). Module Ast-M1 was weakly negatively correlated with total drinking years (R= -0.21) and was positively correlated to liver class (R=0.27) (Fig 5C, Table S8). Module Ast-M1 was enriched in modulation of chemical synaptic transmission (GO:0050804, -log_10_P=59.92), consistent with SNAP-a programs(46) (Fig 5H, Table S10). Ast-M1 was also enriched in immune response categories (R-HSA-1280218, -log_10_P=27.96) (Table S10). Hub genes for module Ast-M1 highlight SNAP-a markers such as *SNAP25* and *SYT1* (Fig 5J, Table S7). Another module of interest was module Ast-M3, 70 genes, which was primarily expressed in cluster Ast2 and positively correlated with total drinking years (R=0.25) (Fig 5B-C, F, Table S7-8). Module Ast-M3 was upregulated in mild and moderate AUD (Fig 5G) and highly enriched in the ECM (M5884, -log_10_P=5.59; R-HSA-1638091, -log_10_P=5.10) (Fig 5I, Table S10). Hub genes included ECM components such as *VCAN*, *TNC* and *GPC6* (Fig 5K, Table S7), reinforcing the role of the extracellular matrix and astrocytes in AUD identified in our differential expression data.

**Figure 5:**
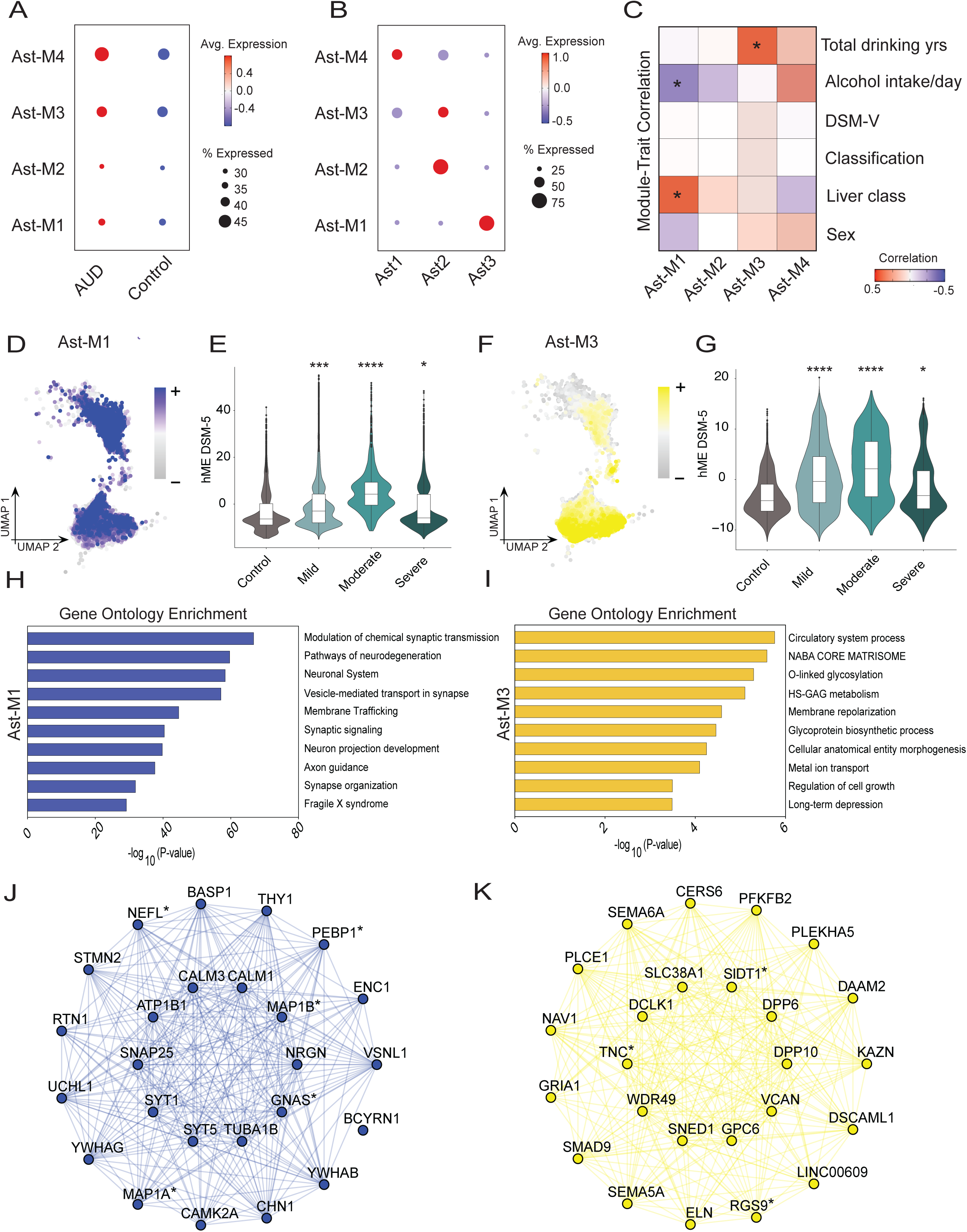
Astrocyte co-expression modules indicate inflammatory phenotype with synapse dysregulation in AUD. (A) Co-expression modules from astrocyte clusters represented by dot plot as the module eigengene (hME) average expression (color of dot) and the percent of genes expressed in each module (size of dot) as a function of (A) AUD classification or (B) cell type cluster. Red indicates increased average expression, blue indicates decreased average expression of the module. (C) Module-trait correlation heatmap for specific traits: Alcohol intake (g/day), DSM-5, Total drinking years, AUD Classification, Liver class and Sex. Color is a function of correlation strength, with red representing positive correlation and blue representing negative correlation with module eigengene expression. *R ≥ |0.2|. (D) UMAP plot colored by modules eigengenes for astrocyte co-expression module Ast-M1. (E) hME expression of astrocyte co-expression module Ast-M1 across DSM-5 categories. (F) Top 10 GO category enrichments for Ast-M1. (G) Visualization of top hub genes for Ast-M1. Genes marked with an asterisk (*) represent hub genes that are also differentially expressed in astrocyte clusters. (H) UMAP plot colored by modules eigengenes for Ast-M3. (I) hME expression of astrocyte co-expression module Ast-M3 across DSM-5 categories. (J) Top 10 GO category enrichments for Ast-M3. (K) Visualization of top hub genes for Ast-M3. For the violin plots in E, I values extend from minimum to maximum, with a white boxplot to show the median value. Color is a function of DSM-5 category. Two-tailed Welch’s t-test for DSM-5 module eigengene expression comparisons, * P<0.05, **P<0.01, ***P<0.001, ****P<0.0001.

The oligodendrocyte network consisted of 5 modules, two of which had increased eigengene expression in AUD (Fig S5A, Table S7). Module Oligo-M2, 427 genes, was primarily expressed in cluster Oligo2, the cluster primarily driving differential expression (Fig S5B, Table S7). Module Oligo-M2 was weakly positively correlated with sex (R=0.19) and upregulated in mild and moderate AUD (Fig S5C-E, Table S8). Module Oligo-M2 was also highly enriched in neurodegenerative pathways (hsa05012, -log_10_P=55.19; WP5124, -log_10_P=10.59) (Fig S5F, TableS10). Several hub genes were related to neurodegenerative disorders such as *CALM1*, *CALM3*, and *SOD1* (Fig S5G, Table S7)(61). Module Oligo-M4 was primarily expressed in cluster Oligo1 and was upregulated in mild and moderate AUD (Fig S5B, H-I). Additionally, module Oligo-M4 was positively correlated with alcohol intake (g/day) (R=0.20) and total drinking years (R=0.25) (Fig S5C, Table S8). Oligo-M4 was enriched in homeostatic oligodendrocyte functions such as myelination (GO:0042552, -log_10_P=17.62) (Fig S5J, Table S10). Hub genes included canonical myelination and oligodendrocyte markers such as *PLP1* and *MAG* (Fig S5K, Table S7).

### Excitatory neuronal modules show modest changes in synaptic organization and **postsynaptic density**

As the second largest class of differentially expressed genes in AUD, we constructed gene co-expression networks for excitatory neurons. Clusters Excit1-7 showed a distinct signature in comparison to clusters Excit8-11 (Fig S6), therefore we chose to construct two separate excitatory neuron networks. Excitatory clusters 8-11 consisted of three gene co-expression modules none of the modules significantly correlated with alcohol traits (Fig S6, Table S7). In contrast, excitatory neurons from cluster Excit1-7 had nine gene co-expression modules, six of which had increased eigengene expression in AUD with one module correlating with total drinking years (R=0.42) (Fig S7A-C, Table S7-8). Excitatory module M3 (Excit-M3), 165 genes, was expressed across clusters Excit3 and Excit7 and was upregulated in moderate and severe AUD (Fig S7B-E, Table S7). Module Excit-M3 was enriched in trans-synaptic signaling (GO:0099537, -log_10_P=9.76), Fragile X syndrome (WP4549, -log_10_P=8.95) as well as alcoholism (hsa05034, -log_10_P=2.47) (Fig S7F, Table S10). Hub genes include genes that have been previously implicated in AUD such as *GRIN2A* and *HOMER1* (Fig S7G)(62).

The inhibitory neuron network consisted of twelve gene co-expression modules, none of which significantly correlated with alcohol traits (Fig S8, Table S8). The modest correlations between alcohol traits and neuronal gene co-expression networks in contrast to the strong correlations observed in glial networks further suggests the importance of glial co-expression networks in AUD and their therapeutic potential.

### GWAS enrichment meta-analysis highlights role for glia in AUD genetic risk

We next used Multi-marker Analysis of GenoMic Annotation (MAGMA) to identify which cell subtypes in this study harbored aggregated genetic risk for AUD using both differentially expressed genes and hdWGCNA modules for gene-set analysis. As expected, specific neuronal subtypes were enriched for SNPs associated with AUD in the million-veteran program GWAS dataset (Fig 6, Table S11)(29). Notably, most cell types enriched for AUD-associated SNPs were glial (Fig 6, Table S11), with the strongest glial enrichments observed for differential expression in microglia and OPCs (Fig 6A-B). Astrocytes and neurons were enriched for AUD risk in the context of specific hdWGCNA modules (Fig 6C). This analysis underscores that changes in expression within these cell types are causal to AUD risk and disease progression.

**Figure 6:**
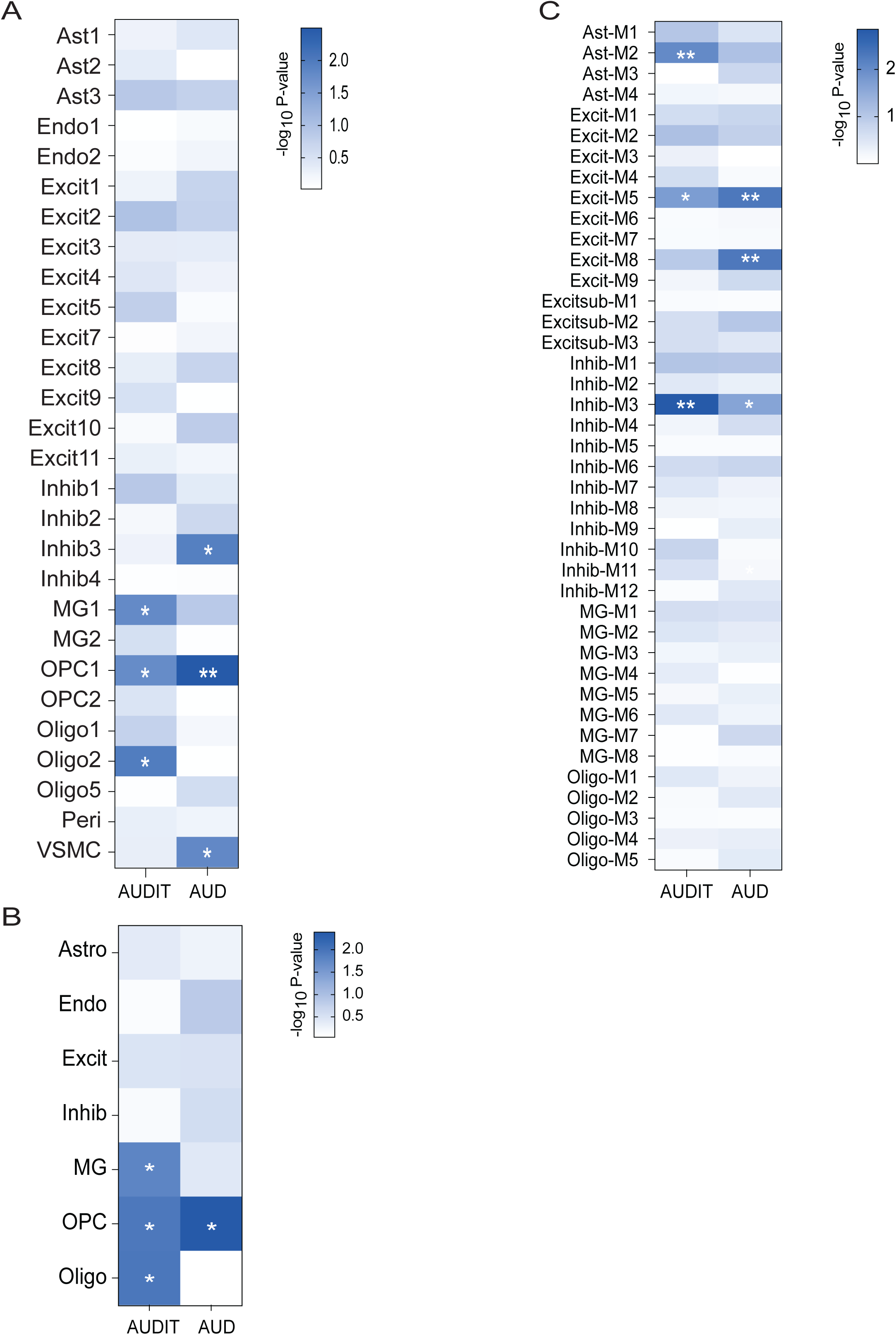
GWAS meta-analysis enrichment for alcohol phenotypes highlights glial risk for AUD disease progression. Heatmaps of GWAS enrichment using MAGMA for alcohol dependence and consumption AUDIT (AUDIT-C). (A) Enrichment of specific cell type clusters for significant differentially expressed genes (DEGs) (B) Enrichment of aggregate cell type classes for significant DEGs. (C). Enrichment of hdWGCNA modules. Color is a function of -log10 (P-value), with dark blue representing stronger enrichment, light blue representing weaker enrichment and white representing no enrichment. *P<0.05, **P<0.01

### Proteome integration replicates broad glial transcriptomic changes observed in AUD

Finally, we integrated our snRNAseq data with a recent proteome study in the dlPFC of AUD individuals(28). Using rank-rank hypergeometric overlap, we determined a significant degree of overlap between concordant differentially abundant protein and DEGs (Fig S9B-C, Table S12) (63). We additionally tested consistency of overlap between the AUD proteome and AUD single cell atlas, which showed consistency between the two datasets on a cell type-specific level (FigS9 D-H, Table S12). Pathway analysis of the overlapping genes/proteins highlighted categories observed in both our dataset and the alcohol field such as the role of the ECM, inflammatory response and dysregulation of neurotransmitter release in AUD (Fig S9I-J, Table S12). As well as additional novel categories such as interferon response, which is consistent with our microglia gene co-expression network analysis (Fig S9H).

## Discussion

The findings presented here substantially refine our understanding of AUD molecular pathology and risk in human brain. Due to the number of samples (n=73) and nuclei (n=459,446), this study was sufficiently powered to capture 32 cell type clusters including novel subtypes of neurons, astrocytes and microglia. Moreover, this study captured significantly more DEGs than any other previous studies in AUD in humans(11,45). There were prominent gene expression changes in astrocytes which identified novel targets in addition to high consistency with previous sequencing(8,11). Furthermore, we are the first to perform WGCNA analysis to identify co-expression networks in individuals with AUD on the single-nuclei level. These co-expression networks identified novel regulatory players that correlated with alcohol specific traits. Our identification of AUD-responsive modules and DEGs highlighted genes that can be either casual or downstream effects of alcohol itself that can be targeted as potential novel therapeutics. A significant overlap of these genes were validated using available proteome data in the PFC of AUD individuals, suggesting these gene expression changes are long-lasting and can affect behavior(28). Finally, the integration of single nuclei clusters with GWAS data for alcohol dependence and AUDIT scores revealed specific glial and neuronal subtypes that are key for AUD risk and disease progression.

Given the complexity of neuropsychiatric disorders such as AUD, disentangling the role of each cell type in the brain is important and requires single-nuclei resolution. For example, the ability to distinguish glial subtypes—including multiple astrocyte, oligodendrocyte and microglia clusters—enabled us to pinpoint changes specific to selective subsets of these glial cells that were alcohol-responsive and potentially linked to the progression of AUD.

In astrocytes, we identified three clusters (Ast1-3), which were alcohol responsive and contained the largest fraction of DEGs compared to any other cell type. These astrocytic clusters fell into three categories: homeostatic, inflammatory and SNAP-a. Changes in homeostatic and inflammatory astrocytes have previously been identified in both our pilot snRNAseq study and mouse models of alcohol dependence(11,64)—this study replicated many of these genes and expanded gene targets due to the size and power of this study. One of the largest categories of DEGs in astrocytes upregulated in AUD was CSPGs and other components of the ECM. CSPG and ECM enrichment was also observed in module Ast-M3, which significantly positively correlated with total drinking years. Local ECM remodeling, dependent on CSPG response to inflammation, has been posited as a crucial component of the altered brain function underlying neuropsychiatric disorders(65,66). Animal models have found CSPGs are elevated after binge drinking(67) and have a regulatory role in aversion-resistant drinking(68) and intermittent drinking(69). Importantly, inflammatory mediators such as interferons and TNFs modulate ECM remodeling, which can impact behavior. This capacity for regulation of behavior may stem from ECM remodeling’s role in regulation of synaptic plasticity, including dendritic spine formation and neurite outgrowth and even myelination(66). Similarly, in the dlPFC from individuals with opioid use disorder, there was significant enrichment in factors that control ECM remodeling, highlighting a common neuroimmune-ECM interaction in substance use disorders(70). Furthermore, proteome integration replicated several key ECM proteins and demonstrated overlapping enrichment of ECM function(28). Taken together this suggests a strong link between neuroinflammation, ECM remodeling and AUD pathology.

In microglia, we identified two clusters that were alcohol responsive and represented loss of homeostatic control leading to an inflammatory phenotype—a common microglial mechanism also observed in other neurological disorders(24,57,71). Beyond a simple dichotomy of homeostatic/inflammatory microglia, due to the sample size and power of this study, we were able to identify novel microglia subtypes in AUD, including border-associated macrophages (BAM) and disease-associated microglia (DAM). BAMs are localized between the brain parenchyma and the periphery, leading to a pivotal role in immune surveillance and inflammatory regulation(20). Recently, BAMs have been implicated in the pathogenesis of neurodegenerative and neuroinflammatory disorders via regulation of immune cell infiltration into the parenchyma(72). Similarly, DAMs have also been implicated in the pathogenesis of multiple neurodegenerative disorders due to their coincident downregulation of homeostatic microglia genes (*P2RY12, CX3CR1, TMEM119*) and upregulation of genes involved in lysosomal, phagocytic and lipid metabolism(22,57). Differentially expressed genes captured both DAM and BAM signatures in addition to inflammatory pathways previously reported in both human studies and mouse models of alcohol dependence such as TLRs, interleukins and cytokine signaling (see (51) for review).

Additionally, microglia modules captured the importance of both canonical immune signaling and these novel microglia/macrophage subtypes. Microglia module MG-M2, which positively correlated with total drinking years and was upregulated in mild/moderate AUD, was highly enriched in loss of homeostatic microglia and upregulation of DAM and interferon-responsive microglia populations. Hub genes in MG-M2 have been shown to regulate alcohol behaviors and synaptic functions. TLR2 decreases alcohol intake in mouse models(73) and IL1B increases GABA release via proinflammatory pathways in alcohol-dependent mice(74). NEAT1, a lncRNA upregulated in alcohol dependent mice and a hub gene in the upregulated microglia module MG-M2, has been shown to mediate and promote inflammation in cell culture and animal models(64,75,76). Proteome integration replicated the importance of DAM and BAM subpopulations as well as inflammatory pathways like interferon signaling in microglia(28). The recruitment of these microglial subpopulations in AUD may have a causative effect since in animal models of alcohol dependence, ablation of microglia blunts escalation of drinking after dependence(38). Moreover, the enrichment of AUD risk in cluster MG1 further causally connects these shifting inflammatory microglial subtypes with regulation of both AUD risk and pathogenesis. However, future studies will be required to link these inflammatory changes to specific microglia subtypes identified in AUD in order to develop microglia subtype-specific treatments.

SPI1 is a key candidate in support of this hypothesis. SPI1/PU.1 is a transcription factor (TF) that has been previously identified as a candidate gene in AUD as well as other neuropsychiatric disorders(77,78). SPI1 is a myeloid lineage determining TF that together with MEF2C shapes the enhancer landscape specific to microglia in brain(79). In human brain, increased SPI1 expression in myeloid lineage cells was associated with a higher risk for AUD and is a candidate gene in AUD(78). Some animal models have also reported that chronic alcohol consumption influences expression of SPI1 in isolated microglia(80). SPI1 also regulates expression of key downstream genes in microglia including homeostatic and DAM genes. We have previously shown in an animal model of dependence that depletion of CSF1R in microglia prevents escalation of alcohol intake(38), suggesting at least one downstream target of SPI1 is a key regulator of alcohol behaviors. Furthermore, increased PU.1 expression in BV2 microglia increased inflammatory response, repressed homeostatic microglial gene expression, and activated the DAM signature(81). These PU.1 regulated functions suggest a risk association of PU.1 to AUD may be driven in part by increased pro-inflammatory response and loss of homeostatic control in specific microglial subtypes(81).

Although most alcohol-responsive populations and DEGs were observed in glial clusters, we ultimately consider these glial effects as a concerted neural response leading to neuronal changes that will alter behavior. These changes likely occur both via alcohol’s direct interactions with neurons and the neuroinflammatory effect of alcohol on glial populations that alters neuronal firing (see(51,62) for review). We identified multiple excitatory and inhibitory subtypes with layer- and peptide-specific markers that were alcohol responsive. Inhibitory and excitatory neurons represented 25% of DEGs and the categories enriched in these subclusters were canonical to AUD, including changes in glutamatergic and GABAergic signaling as well as novel links to Fragile X Syndrome. One previous study linked ethanol-dependent changes in GABA_B_R expression and dendritic plasticity to fragile-X mental retardation protein (FMRP) and even found lack of these ethanol-dependent changes in FMR1 knockout mice, suggesting FMRP is an important regulator of protein synthesis and synaptic plasticity following alcohol exposure(82). Additionally, genetic risk for AUD was enriched in inhibitory clusters and modules as well as the excitatory modules mentioned above. How these neuronal firing changes are linked with neuroinflammatory mechanisms that incur AUD risk and influence pathology will be investigated in the future.

Our study is not without limitations. Almost all individuals were of European ancestry (n=71), limiting the generalizability of our results. Most individuals included in our study were also male (N=54) with a small cohort of female individuals (N=19). The general linearized model with sex as a variable showed no significant effect, allowing us to pool samples. We additionally included sex as a covariate in the differential expression analysis as to not bias our results. Five out of the 41 hdWGCNA modules had significant correlations with sex, suggesting sex does influence co-expression networks in AUD, but is likely underpowered in our study. Future studies will investigate sex-specific differences in AUD. Technical limitations with snRNA-seq of human brain samples have been previously described(83). We, like others, found a much greater proportion of neurons compared to glial cells but due to the sample size and extraction technique were able to capture more non-neuronal cells than typically reported. Recent studies reported between 1.6-2.1% microglia(32,84); whereas, we captured 4.9% microglia, allowing us to report on microglia subpopulations in AUD for the first time. Moreover, despite droplet-based snRNAseq not capturing genes expressed at very low levels, we were still able to identify differential gene expression for thousands of genes in defined cell types.

To date, this is the largest and most well-powered single-nucleus transcriptomic atlas of the human brain in AUD. Our data illustrates the utility of this type of complex, single cell data in understanding the molecular etiology of AUD pathology and risk. Our database of single-nuclei analysis in the adult brain is also made available with this study and provides a resource to elucidate the biological mechanisms underlying AUD. We have demonstrated individuals with AUD have broad transcriptomic dysregulation in multiple cell types, specifically in glia. Moreover, our results show a specific role for novel glial subtypes, especially DAM/BAM microglia and SNAP-a astrocytes, underlying genetic risk of AUD. We expect that this dataset will be crucial for designing informed experiments to identify the biological mechanisms and novel drug targets in AUD.

## Acknowledgements

The authors thank The University of Texas at Austin’s Genomic Sequencing and Analysis Facility (GSAF) and the Ichan School of Medicine at Mount Sinai Genomic Core Facility and its members for preparing and sequencing the cDNA libraries. We are also grateful to the New South Wales Tissue Resource Centre at the University of Sydney for providing human brain samples.

## Conflict of interest

The authors declare no competing interests.

## Supplementary Materials

Figure S1: *Metadata and sequencing QC information*

(A) Ncells vs. NGenes between sequencing sites as a function of log_10_(nFeature). (B) Ncells between sequencing sites shows most cells are from MSSM but both sequencing sites are equally proportioned between control and AUD individuals. (C) Cell density distribution of nCount between sequencing sites before trimming. (D) nFeature vs. nCount correlation between sequencing sites as a function of percent mitochondrial contamination. Frequency boxplots for metadata categorical variables (E) Sex (F) AUD Classification (G) Smoking frequency (H) Ethnicity (I) DSM-5 categories (J) Liver class. Density plots for continuous metadata variables (K) Pack years (1 packet/day/1 yr) (L) Age (M) Postmortem interval (PMI) (N) Alcohol Use Disorders Identification Test (AUDIT) (O) Total drinking years (yrs) (P) Alcohol intake (g/day).

Figure S2: *Benchmarking of integration of two sequencing sites datasets using Harmony*

UMAP plots of sequencing site contributions to unsupervised clustering (A) before Harmony integration and (B) after Harmony integration across the final 32 clusters. PCA of individual samples (C) before Harmony integration and (D) after Harmony integration (C=control, A=AUD). UMAP plots of metadata variable contribution to unsupervised clustering after Harmony integration, (E) Liver class, (F) Smoking frequency, (G) AUD classification, and (H) Sex.

Figure S3: *Aggregate cell type proportions for nuclei between metadata conditions*

Cell type proportions for nuclei by cluster for the following metadata: (A) Alcohol intake (g/day), (B) AUD Classification, (C) Total drinking yrs, (D) DSM-5 category, (E) Sample Number (#), (F) Sex. Color represents individual cell type specific clusters (32 final clusters). Y-axis represents the percent of cells as a fraction. X-axis is either a quantitative measure or a categorical variable.

Figure S4: *Astrocyte subtype clustering*

(A) Astrocyte subtype markers to differentiate clusters Ast1-3. Common astrocyte markers include (SLC1A2, SLC1A3, NDRG2, AQP4, GFAP), which are expressed across all three clusters. Ast1-2 specific makers (POU2F1, GPC5, ALDH1L1). Ast2 specific inflammatory markers (CD44, S100B). Synaptic markers highly expressed in Ast3 in comparison to Ast1-2 clusters (GRIN2A, GRIN2B, NPTXR, DNM1, YWHAG, CAMK2A, RTN4). (B) Circle plot showing percentage of nuclei in each astrocyte cluster contributing to the total number of astrocyte nuclei (n=46,247). (C) Feature Plots showing layer specific astrocyte makers across isolated UMAP for astrocyte clusters Ast1-3. Layer1-6 markers (FGFR3, SLC14A1) and Layer1-2 markers (GFAP, SLC1A3). Ast1-2 express both types of astrocytic depth markers, whereas Ast3 expresses only Layer1-2 markers. Color is a function of average expression, with red indicating high expression and grey representing no expression. Top 20 GO category enrichments for (D) Ast1, (E) Ast2, (F) Ast3. Enrichment is a function of -log10 P-value. (G) Overlap of common GO categories between astrocyte clusters. (H) Number of gene overlaps with synaptic neuron and astrocyte program astrocytes (SNAP-a) GO categories for each cluster. Ast3 had the highest gene overlaps with SNAP-a categories. (I) UMAP plot of SNAP-a gene set expression in astrocyte clusters (Ast1-3). Ast3 had high concerted expression of the SNAP-a signature suggesting a functional difference in astrocytic function than Ast1-2. Color indicates average expression of SNAP-a gene expression signature, with yellow indicating high expression, green indicating low expression and purple indicating no expression. For the violin plots in A, values extend from minimum to maximum, color is used to differentiate between markers.

Figure S5: *Oligodendrocyte co-expression modules highlight myelination and lipid metabolism roles in AUD*

(A) Five co-expression modules from oligodendrocyte clusters represented by dot plot as the module eigengene (hME) average expression (color of dot) and the percent of genes expressed in each module (size of dot) as a function of (A) AUD classification or (B) cell type cluster. Red indicates increased average expression, blue indicates decreased average expression of the module. (C) Module-trait correlation heatmap for specific traits: Alcohol intake (g/day), DSM-5, Total drinking years, AUD Classification, Liver class and Sex. Color is a function of correlation strength, with red representing positive correlation and blue representing negative correlation with module eigengene expression. *R ≥ |0.2|. (D) UMAP plot colored by modules eigengenes for oligodendrocyte co-expression module Oligo-M2. (E) hME expression of oligodendrocyte co-expression module Oligo-M2 across DSM-5 categories. (F) Top 10 GO category enrichments for Oligo-M2. (G) Visualization of top hub genes for Oligo-M2. (H) UMAP plot colored by modules eigengenes for Oligo-M4. (I) hME expression of oligodendrocyte co-expression module Oligo-M4 across DSM-5 categories. (J) Top 10 GO category enrichments for Oligo-M4. (K) Visualization of top hub genes for Oligo-M4. For the violin plots in E, I values extend from minimum to maximum, with a white boxplot to show the median value. Color is a function of DSM-5 category. Two-tailed Welch’s t-test for DSM-5 module eigengene expression comparisons, * P<0.05, **P<0.01, ***P<0.001, ****P<0.0001.

Figure S6: *hdWGCNA analysis of excitatory subclusters (Excit8-11)*

(A) PCA plot of excitatory neuron clusters (Excit1-11), showing separation of clusters Excit8-11. (B) Network analysis was performed on these subclusters resulting in three co-expression modules represented by dot plot as the module eigengene (hME) average expression (color of dot) and the percent of genes expressed in each module (size of dot) as a function of (B) AUD classification or (C) cell type cluster. Red indicates increased average expression, blue indicates decreased average expression of the module. (D) Module-trait correlation heatmap for specific traits: Alcohol intake (g/day), DSM-5, Total drinking years, AUD Classification, Liver class and Sex. Color is a function of correlation strength, with red representing positive correlation and blue representing negative correlation with module eigengene expression. No co-expression modules significantly correlated with the selected traits.

Figure S7: *hdWGCNA analysis of excitatory subclusters (Excit1-7)*

(A) Nine co-expression modules from excitatory neuron clusters (Excit1-7) represented by dot plot as the module eigengene (hME) average expression (color of dot) and the percent of genes expressed in each module (size of dot) as a function of (A) AUD classification or (B) cell type cluster. Red indicates increased average expression, blue indicates decreased average expression of the module. (C) Module-trait correlation heatmap for specific traits: Alcohol intake (g/day), DSM-5, Total drinking years, AUD Classification, Liver class and Sex. Color is a function of correlation strength, with red representing positive correlation and blue representing negative correlation with module eigengene expression. *R ≥ |0.2|. (D) UMAP plot colored by modules eigengenes for excitatory neuron co-expression module Excit-M3. (E) hME expression of excitatory neuron co-expression module Excit-M3 across DSM-5 categories. (F) Top 10 GO category enrichments for Excit-M3. (G) Visualization of top hub genes for Excit-M3. Genes marked with an asterisk (*) represent hub genes that are also differentially expressed in excitatory neuron (Excit1-7) clusters For the violin plots in E values extend from minimum to maximum, with a white boxplot to show the median value. Color is a function of DSM-5 category. Two-tailed Welch’s t-test for DSM-5 module eigengene expression comparisons, * P<0.05, **P<0.01, ***P<0.001, ****P<0.0001.

Figure S8: *hdWGCNA analysis of inhibitory neurons*

Twelve co-expression modules from inhibitory neurons represented by dot plot as the module eigengene (hME) average expression (color of dot) and the percent of genes expressed in each module (size of dot) as a function of (A) AUD classification or (B) cell type cluster. Red indicates increased average expression, blue indicates decreased average expression of the module. (C) Module-trait correlation heatmap for specific traits: Alcohol intake (g/day), DSM-5, Total drinking years, AUD Classification, Liver class and Sex. Color is a function of correlation strength, with red representing positive correlation and blue representing negative correlation with module eigengene expression. No co-expression modules significantly correlated with the selected traits.

Figure S9: *Significant overlaps between single nuclei RNAseq and proteome in cortex of AUD individuals*

(A) Rank-rank hypergeometric overlap heatmap. Each pixel represents the overlap between this study’s snRNA-seq differentially expressed genes (DEGs) and the Teng et al., proteome differentially abundant proteins (DAPs). The significance of overlap is a function of -log10P, color coded with a step size of 200. High enrichment in the bottom left corner denotes co-upregulate between studies. (B) Venn diagram of overlap of most co-downregulated or (C) co-upregulated genes and proteins. (D) Global consistency analysis of snRNA-seq versus proteome. Agreement or disagreement was estimated by deviation from random expectation (*z*-score) of single-cell DEG average rank scores in the ranked list of DAPs from proteome differential analysis. A negative z-score denotes consistency, a positive z-score suggests disagreement between datasets. (E) Heatmap of selected overlaps between snRNAseq and protein. Color is a function of log2 fold change, with red representing upregulation and blue representing downregulation. Cluster identification for each gene/protein is color coded as a separate column to the left of the heatmap. Violin plots of selected genes/proteins between Control and AUD individuals: (F) CD44, (G) MCU, (H) DCKL1. Top row represents our snRNAseq experiment. Bottom row represents protein. (H) Ingenuity Pathway Analysis (IPA) for overlapping genes and proteins. (I) Most co-upregulated genes/protein pathway enrichment analysis. (J) Most co-downregulated genes/protein pathway enrichment analysis. Enrichment is represented as a function of -log10 (P-value). For the violin plots in F-H values extend from minimum to maximum, with a boxplot to show the median value. For the snRNAseq expression (top row), dots represent outliers. For the protein level (bottom row), dots represent individual samples. Graphs for specific proteins were obtained from (https://www.lmdomics.org/AUDBrainProteomeAtlas/) . Color is a function of AUD classification category. * P<0.05, **P<0.01.

Table S1: *Cohort metadata and sequencing QC per cohort*

Table S2: *Correlation coefficients for different generalized linear models between metadata variables after thresholding and normalization and harmony integration.* (A) nFeatures (B) nGenes

Table S3: *Cell type proportion statistical testing for glial clusters MG1-2, Ast1-3, Oligo1-5*.

Table S4: *GO enrichment & pathway analysis for glial clusters of interest*

Table S5: *Differentially expressed genes in AUD*

Table S6: *GO enrichment & pathway analysis for differentially expressed genes in AUD*

TableS7: *hdWGCNA modules for each cell type*

Table S8: *hdWGCNA module-trait correlations for each cell type*

Table S9: *Collated external gene sets*

TableS10: *GO enrichment & pathway analysis for WGCNA modules*

Table S11: *GWAS MAGMA analysis statistics*

Table S12: *Proteome overlap*

## Notes

### Competing Interest Statement

The authors have declared no competing interest.

### Summary of Updates

The revised manuscript contains updated methods.

