## SupplementaryMaterials for "Integrative genomics approach identifies glial transcriptomic dysregulation and risk in the cortex of individuals with Alcohol Use Disorder"

#### Supplemental Methods

##### Cohort selection and postmortem tissue collection

All donors were required to meet the following criteria to be considered in this study: no head injury at time of death, lack of developmental disorder, no recent cerebral stroke, no history of other psychiatric or neurological disorders, no history of intravenous or polydrug abuse, no severe cirrhosis. Diagnosis of alcohol dependence as based on DSM-5 criteria. The postmortem interval (PMI) represents the delay between an individual's death and collection and processing of the brain. To assess RNA quality, we measured the RNA integrity number (RIN) obtained for our samples. AUD and control donors were selected to be as similar as possible in terms of age, sex, PMI, pH of tissue, and cause of death (Table S1).

##### Droplet-based single-nucleus RNA-seq

For droplet-based single-nucleus RNA sequencing (snRNA-seq), libraries were prepared as previously described (1) using the Chromium Single Cell 3' Reagent Kits v3.1 according to the manufacturer's protocol (10X Genomics, Pleasanton, CA) with a target cell count of at least 10,000 cells. The generated snRNA-seq libraries were sequenced using NovaSeq 6000 using a S4 flow cell. Of the 73 total samples, 50 samples were processed at Icahn School of Medicine at Mount Sinai and 23 samples were processed at the University of Texas at Austin (Table S1). Sequencing site was accounted for in all downstream analyses.

Fastq files were processed using the Cellranger mkfastq command (v 6.1.2) and default parameters starting with the refdata-cellranger-GRCh38-1.2.0 transcriptome as per the instructions provided by 10x Genomics. Reads were demultiplexed by the sample index using the cellranger. Fastq files were aligned to the custom transcriptome, cell barcodes were demultiplexed and UMIs corresponding to genes were counted using the cellranger count command with default parameters.

##### Filtering and normalization

Ambient RNA was corrected using SoupX (v1.3.0) as previously described(2). The corrected matrix was then loaded into Seurat (v 4.2.0). Based on the distribution of nGene (total number of genes detected in each cell) for the total dataset, barcodes associated with fewer than 200 detected genes were removed. Based on the distribution of nUMI (the total number of UMIs detected in each cell), the top quartile of barcodes was also excluded as most likely being multiplets and to minimize our sequencing site batch effect. Nuclei with more than 20% of UMI counts attributed to mitochondrial genes were also removed. Our custom filtering helped increase the number of glia cells recovered (data not shown).

The UMI counts were normalized to 10,000 counts per cell and converted to log scale (Seurat function NormalizeData). Sample data from Table S1 was added to the object as metadata; mitochondrial reads were regressed out to minimize downstream effect. The Seurat FindVariableGenes function was used with default cut-offs (nfeatures=2000). The highly variable genes were used to calculate 40 PCs. Based on the PC elbow plot of the standard deviation of the PCs, the first twenty PCs were retained for downstream analysis. The FindNeighbors,

FindClusters and RunUMAP functions were applied with a resolution of 0.5, which produced 30 initial clusters.

**Filtering and normalization**

Based on the distribution of nGene (total number of genes detected in each cell) for the total dataset, barcodes associated with fewer than 200 detected genes were removed. Based on the distribution of nUMI (the total number of UMIs detected in each cell), the top quartile of barcodes was also excluded as most likely being multiplets and to minimize our sequencing site batch effect. Nuclei with more than 20% of UMI counts attributed to mitochondrial genes were also removed. Our custom filtering helped increase the number of glia cells recovered (data not shown).

The UMI counts were normalized to 10,000 counts per cell and converted to log scale (Seurat function NormalizeData). Sample data from Table S1 was added to the object as metadata; mitochondrial reads were regressed out to minimize downstream effect. The Seurat FindVariableGenes function was used with default cut-offs (nfeatures=2000). The highly variable genes were used to calculate 40 PCs. Based on the PC elbow plot of the standard deviation of the PCs, the first twenty PCs were retained for downstream analysis. The FindNeighbors, FindClusters and RunUMAP functions were applied with a resolution of 0.5, which produced 30 initial clusters.

**Integrating cohorts between sequencing sites**

Due to batch effects between sequencing sites, we next integrated our cohort using Harmony (v1.1.0)(3). Harmony is a general-purpose R package with an efficient algorithm for integrating multiple data sets. It is especially useful for large single-cell datasets such as single-cell RNA-seq. After successful integration with Harmony, clusters were visualized using UMAP to ensure no batch effects remained (Figure S2). We additionally ran a regression model with the metadata to determine if sequencing site batch was removed and if any additional metadata needed correction (Table S2). No metadata significantly correlated with nUMI, nGene or percentage mitochondria marked genes (Table S2), signifying harmony corrected our batch effect. After harmony correction, we observed 30 initial clusters.

**Marker identification for cell type specific clusters**

Although clusters obtained using Seurat default settings may correspond to known biological cell types, some clusters may appear to potentially identify entirely new cell types or splinter existing cell types into multiple subtypes. Deciding how and which cell types may come from each cluster can be difficult and at times impossible from available data due to the nature of single nuclei sequencing.

To address this issue, we used tools in the Seurat package to initially determine cluster differences/similarities using hierarchical clustering of the graph-based clusters (BuildClusterTree). Next, we assessed the nodes of dendrogram using a random forest classifier (AssessNodes). Next we characterized clusters for a set of marker genes defined by differential expression analysis of the cells group in each cluster against the remaining cells within the corresponding aggregate cell type cluster using Seurat FindMarkers function. This analysis was applied to all clusters independently. Significantly over-expressed genes were defined based on the Wilcoxon-rank-sum test with an FDR <0.05 and an absolute value log2FC of 0.5. Only genes detected in at least 25% of the cells within the given subcluster were considered. Gene ontology (GO) enrichment analysis was performed using Metascape(4). Reference datasets were used to verify cell-type ontologies (5–12) From this combined analysis we identified several clusters that required further verification. Three clusters had dual cell-type identification and in the top 0.5% of UMI counts, suggesting multiplets. These were removed from downstream analysis. An additional cluster contained excitatory neuron markers as well as astrocytic markers and a range of UMI counts, suggesting subclustering was required. We subclustered at a resolution of 0.5 to produce five separate clusters that were reintegrated into the original Seurat object using the Seurat function FindSubCluster. Four of these five subclusters were labeled as excitatory neurons (Excit8-11) and one was labeled as astrocytes (Ast3) with no dual cell type identification. The proximity of this astrocytic cluster to excitatory neuron clusters was investigated further in Fig S4. Re-running BuildClusterTree and a Spearman correlation of average cluster expression determined the hierarchical relationship was related to other similarly identified cell types (Fig 1). This resulted in 32 final clusters for downstream analysis.

**Consistency analysis**

Consistency of gene-expression perturbations in the different cell types observed in snRNAseq data with those detected in AUD bulk RNA-seq or proteome was assessed using a resampling approach. To test whether the genes identified as DEGs were also detected as high ranking in the differential analysis in bulk or protein data, a *z*-score statistic was computed to quantify the deviation of the observed differential (*P* value) rank scores obtained in the bulk or proteome analysis for the genes detected as DEGs in single cells. This analysis was performed for each cell type independently for both AUD bulk RNA(13) and proteomic data (14). Rank-rank hypergeometric overlap (RRHO) was additionally performed for proteomic integration to determine the significance of overlap between datasets, as previously described (15,16).

**External data sources**

Markers were obtained from the following literature: disease-associated microglia (DAM) (11,12), border-associated microglia (BAM) (17), white matter-associated microglia (WAM) (18), lipid droplet accumulating microglia (LDAM) (19), activated response microglia (ARM) (20), interferon response microglia (IRM) (11), homeostatic microglia (21), proliferative-region-associated microglia (PAM) (22), antigen-presenting microglia (HLA) (11) and inflammatory astrocytes (7,9). Oligodendrocyte subtype markers were collected from (5,6).

**Cluster Annotation**

Genes used as markers for neuron or layer specificity are listed below. The following excitatory neuron layer-specific markers were used: layer II: GLRA3; layer II-III: UNC5D and FGF13; layers II–IV: CUX2, LINC01378, RASGRF2; layer III-V: MAP1A, TPPP, UBB, FTH1; layer IV/V: RORB; layers IV–VI: GRIK4; layer V-VI: TOX, RXFP1, FOXP2, layer VI: COL24A1 SYNPR.

Inhibitory neuron subtypes were annotated based on the expression of canonical inhibitory interneuron markers SST, VIP, LAMP5, CCK and RELN.

The following markers were used for major cell types: macrophage/microglia: CSF1R, P2RY12, APBB1IP; astrocytes: GFAP, ALDH1L1, GJA1, AQP4; OPCs: VCAN, PCDH15; oligodendrocytes: PLP1, MAG; endothelial cells: FLT1, VWF, EPAS1; vascular smooth muscle cells: ACTA2; pericytes: DCN, PDGFRB; excitatory neurons: SLC17A7, SATB2, NPTX1, CDH22; inhibitory neurons: GAD1, GAD2 as previously described (8,10,15,23,24). Microglia subtype markers based on current literature can be found in Supplemental Table 9 (Table S9). Astrocytic subtype markers for the SNAP-a program was defined as recently reported (24).

Fig S1

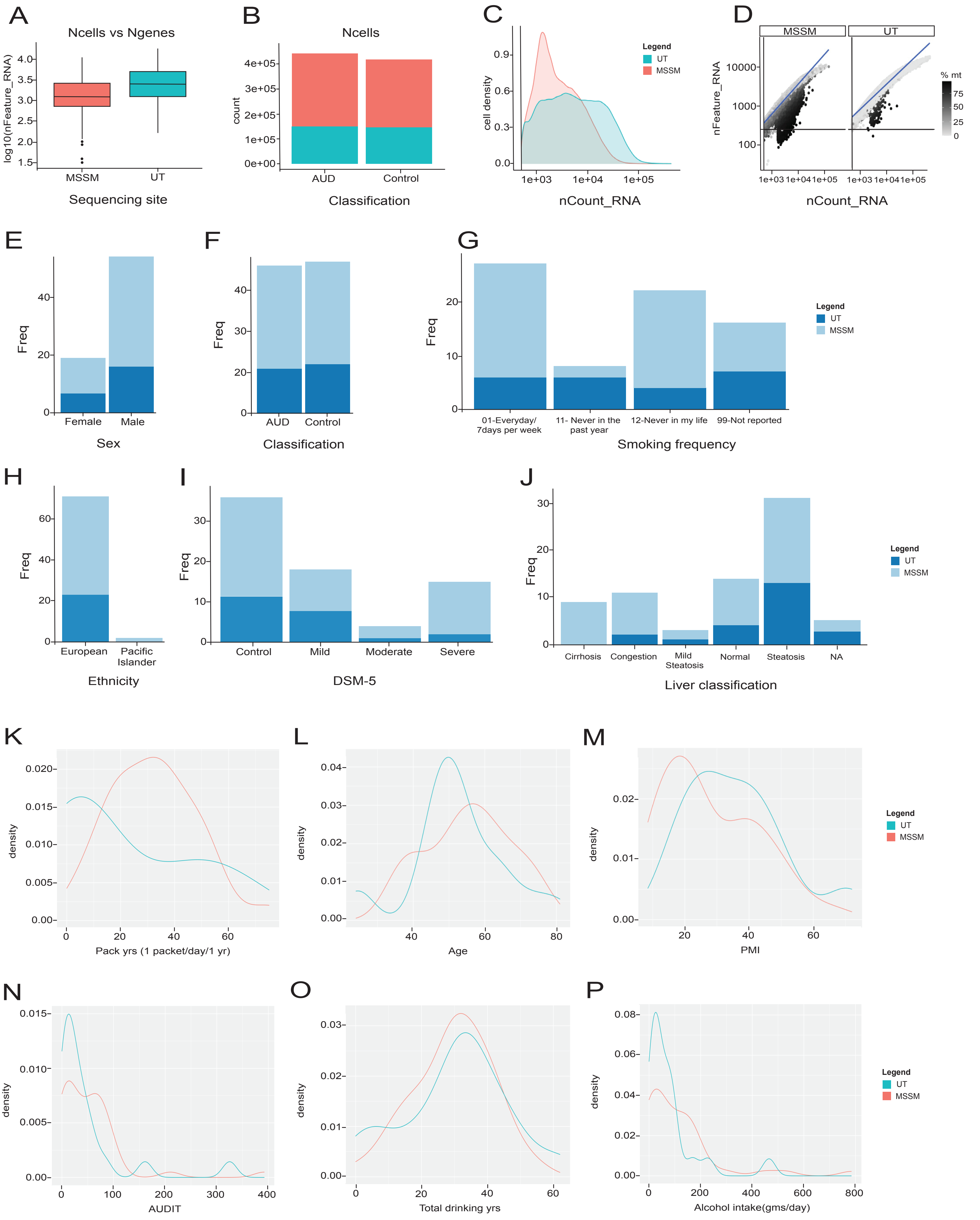
A
Ncells vs Ngenes
log10(nFeature_RNA)
MSSM
UT
Sequencing site
B
Ncells
count
AUD
Control
Classification
C
cell density
Legend
UT
MSSM
nCount_RNA
D
MSSM
UT
nFeature_RNA
% mt
nCount_RNA
E
Freq
Female
Male
Sex
F
Freq
AUD
Control
Classification
G
Freq
01-Everyday/
7days per week
11- Never in the
past year
12-Never in my life
99-Not reported
Smoking frequency
Legend
UT
MSSM
H
Freq
European
Pacific
Islander
Ethnicity
I
Freq
Control
Mild
Moderate
Severe
DSM-5
J
Freq
Cirrhosis
Congestion
Mild
Steatosis
Normal
Steatosis
NA
Liver classification
Legend
UT
MSSM
K
density
Pack yrs (1 packet/day/1 yr)
L
density
Age
M
density
PMI
Legend
UT
MSSM
N
density
AUDIT
O
density
Total drinking yrs
P
density
Alcohol intake(gms/day)
Legend
UT
MSSM

Fig S2

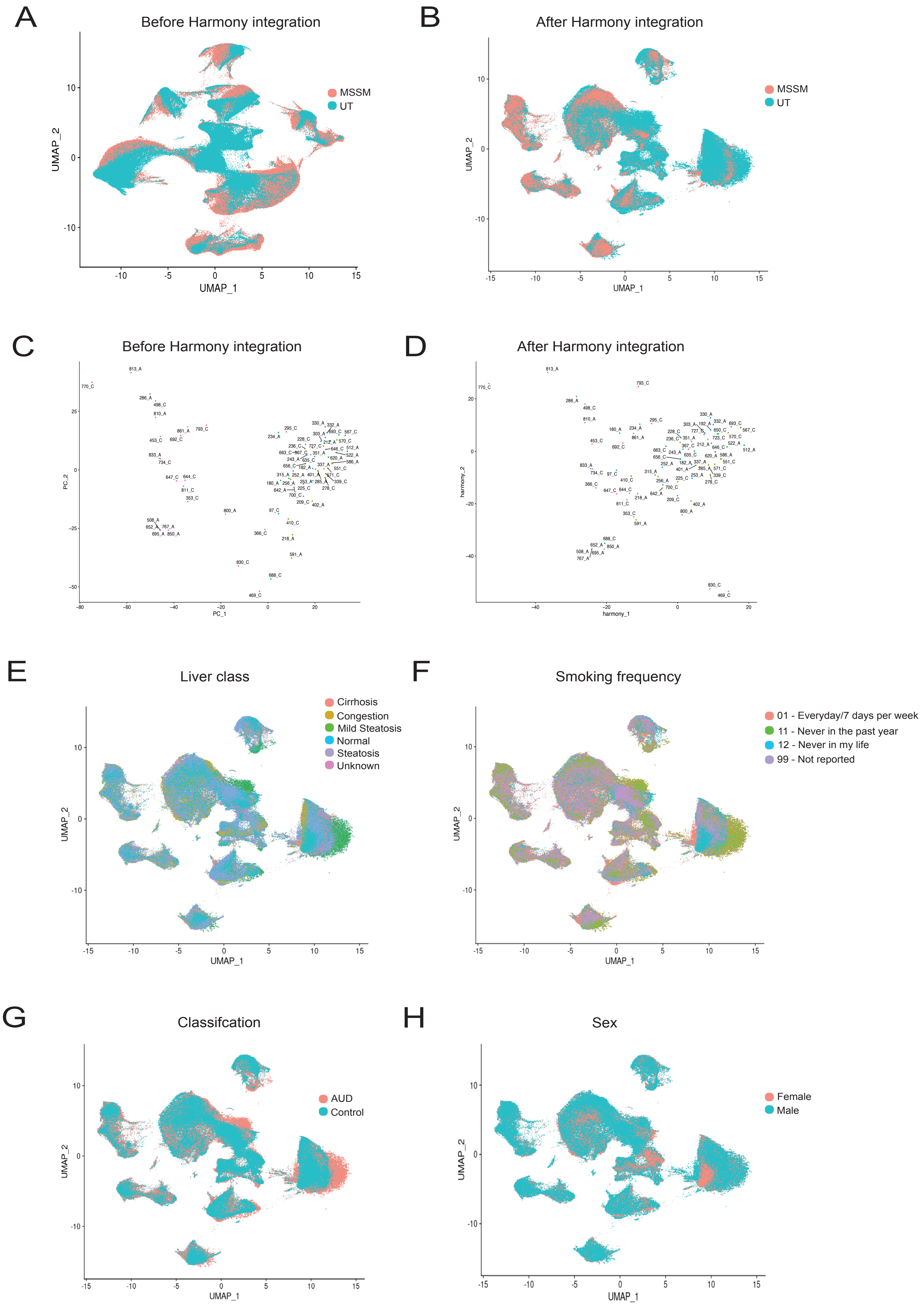
A
Before Harmony integration
MSSM
UT
UMAP_1
UMAP_2
B
After Harmony integration
MSSM
UT
UMAP_1
UMAP_2
C
Before Harmony integration
PC_1
PC_2
D
After Harmony integration
harmony_1
harmony_2
E
Liver class
Cirrhosis
Congestion
Mild Steatosis
Normal
Steatosis
Unknown
UMAP_1
UMAP_2
F
Smoking frequency
01 - Everyday/7 days per week
11 - Never in the past year
12 - Never in my life
99 - Not reported
UMAP_1
UMAP_2
G
Classifcation
AUD
Control
UMAP_1
UMAP_2
H
Sex
Female
Male
UMAP_1
UMAP_2

FigS3

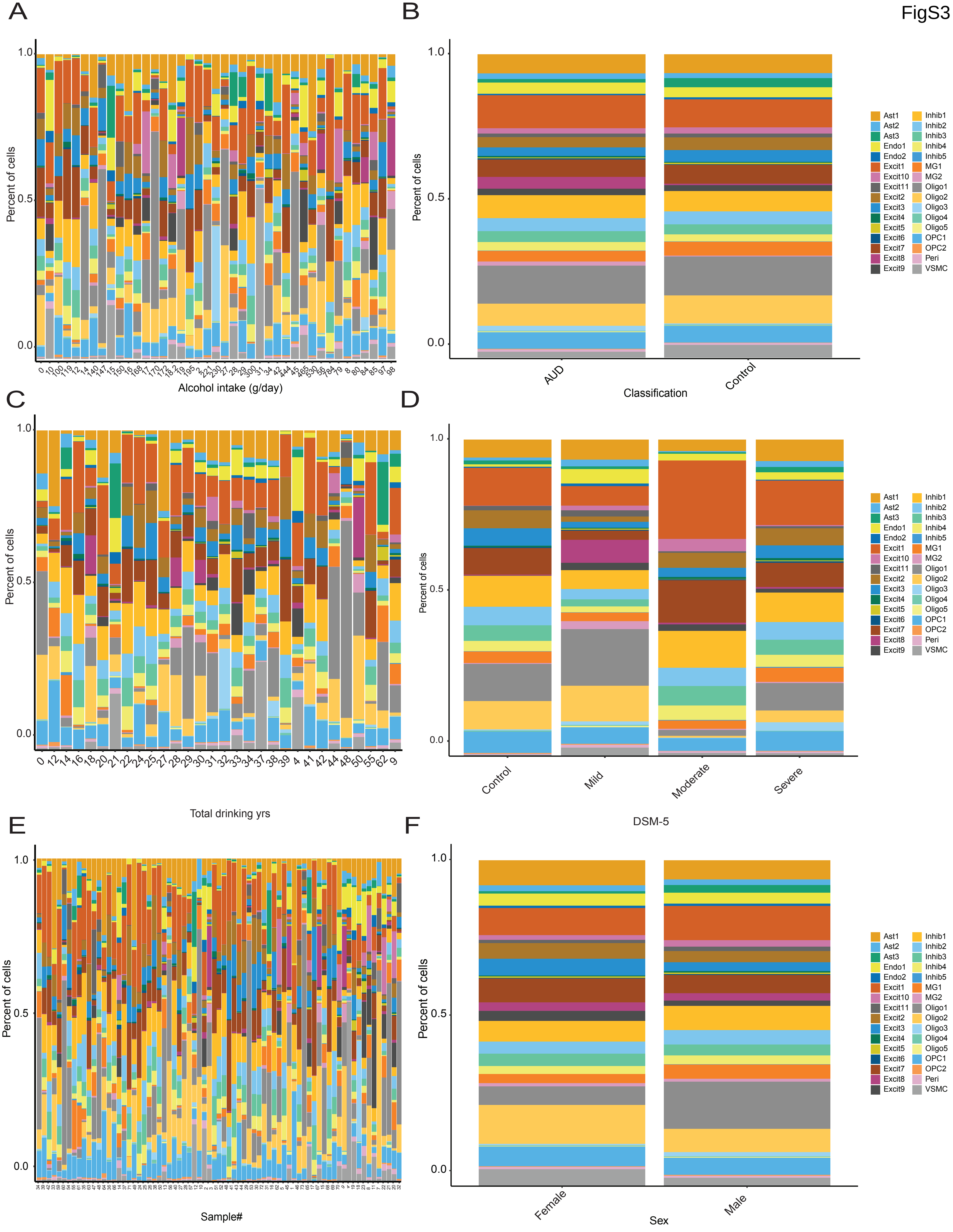

A
Percent of cells
1.0
0.5
0.0
0 10 100 119 12 14 140 147 15 150 16 168 17 170 172 182 19 195 2 221 230 27 28 29 300 31 34 42 444 45 465 530 56 784 79 8 80 84 85 97 98
Alcohol intake (g/day)
B
AUD
Control
Classification
Ast1 Ast2 Ast3 Endo1 Endo2 Excit1 Excit10 Excit11 Excit2 Excit3 Excit4 Excit5 Excit6 Excit7 Excit8 Excit9
Inhib1 Inhib2 Inhib3 Inhib4 Inhib5 MG1 MG2 Oligo1 Oligo2 Oligo3 Oligo4 Oligo5 OPC1 OPC2 Peri VSMC
C
0 12 14 16 18 20 21 22 24 25 27 28 29 30 31 32 33 34 37 38 39 4 41 42 44 48 50 55 62 9
Total drinking yrs
D
Control
Mild
Moderate
Severe
DSM-5
E
Sample#
F
Female
Male
Sex

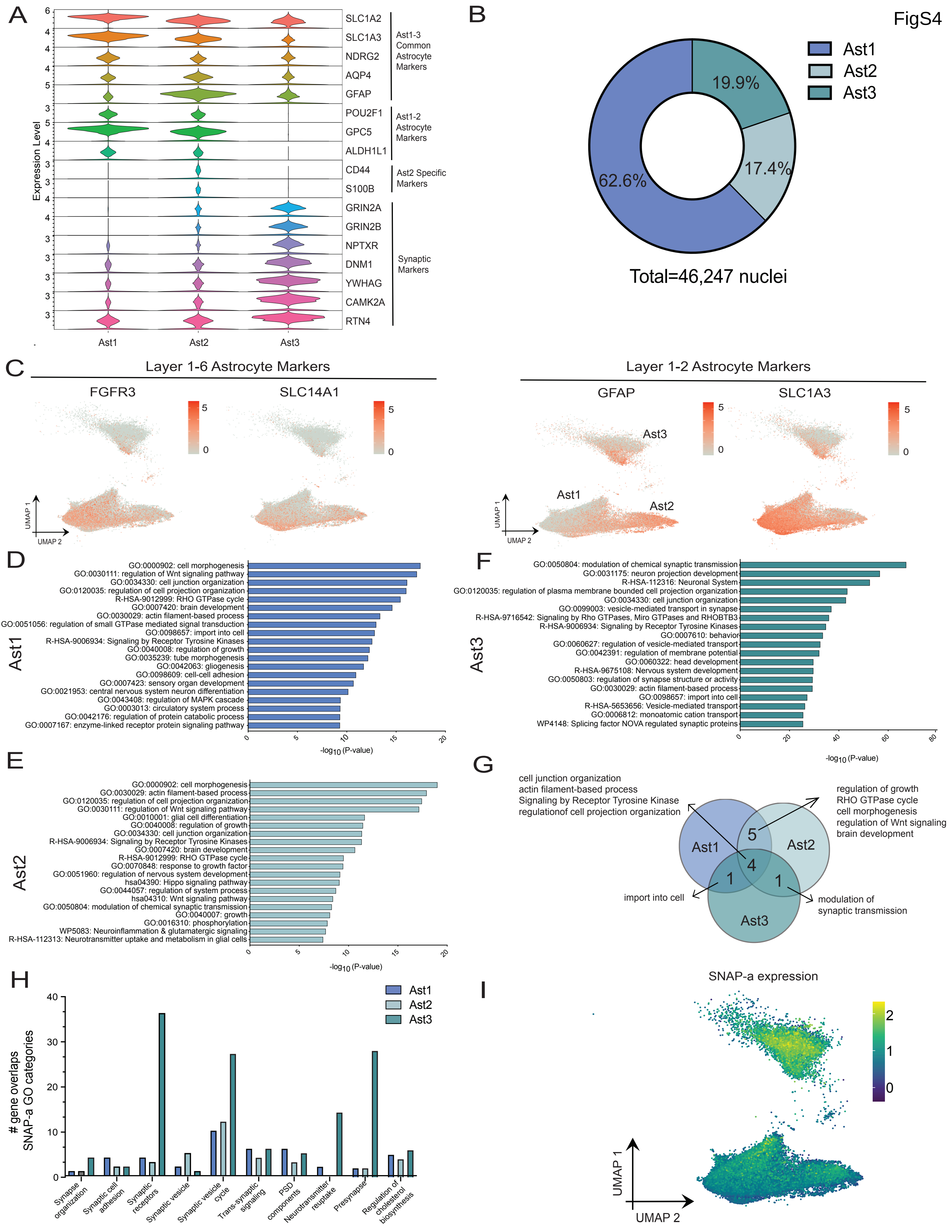
FigS4
A
Expression Level
SLC1A2
SLC1A3
NDRG2
AQP4
GFAP
Ast1-3 Common Astrocyte Markers
POU2F1
GPC5
ALDH1L1
Ast1-2 Astrocyte Markers
CD44
S100B
Ast2 Specific Markers
GRIN2A
GRIN2B
NPTXR
DNM1
YWHAG
CAMK2A
RTN4
Synaptic Markers
Ast1
Ast2
Ast3
B
Ast1
Ast2
Ast3
19.9%
17.4%
62.6%
Total=46,247 nuclei
C
Layer 1-6 Astrocyte Markers
FGFR3
SLC14A1
Layer 1-2 Astrocyte Markers
GFAP
SLC1A3
UMAP 1
UMAP 2
D
Ast1
GO:0000902: cell morphogenesis
GO:0030111: regulation of Wnt signaling pathway
GO:0034330: cell junction organization
GO:0120035: regulation of cell projection organization
R-HSA-9012999: RHO GTPase cycle
GO:0007420: brain development
GO:0030029: actin filament-based process
GO:0051056: regulation of small GTPase mediated signal transduction
GO:0098657: import into cell
R-HSA-9006934: Signaling by Receptor Tyrosine Kinases
GO:0040008: regulation of growth
GO:0035239: tube morphogenesis
GO:0042063: gliogenesis
GO:0098609: cell-cell adhesion
GO:0007423: sensory organ development
GO:0021953: central nervous system neuron differentiation
GO:0043408: regulation of MAPK cascade
GO:0003013: circulatory system process
GO:0042176: regulation of protein catabolic process
GO:0007167: enzyme-linked receptor protein signaling pathway
-log10 (P-value)
E
Ast2
GO:0000902: cell morphogenesis
GO:0030029: actin filament-based process
GO:0120035: regulation of cell projection organization
GO:0030111: regulation of Wnt signaling pathway
GO:0010001: glial cell differentiation
GO:0040008: regulation of growth
GO:0034330: cell junction organization
R-HSA-9006934: Signaling by Receptor Tyrosine Kinases
GO:0007420: brain development
R-HSA-9012999: RHO GTPase cycle
GO:0070848: response to growth factor
GO:0051960: regulation of nervous system development
hsa04390: Hippo signaling pathway
GO:0044057: regulation of system process
hsa04310: Wnt signaling pathway
GO:0050804: modulation of chemical synaptic transmission
GO:0040007: growth
GO:0016310: phosphorylation
WP5083: Neuroinflammation & glutamatergic signaling
R-HSA-112313: Neurotransmitter uptake and metabolism in glial cells
-log10 (P-value)
F
Ast3
GO:0050804: modulation of chemical synaptic transmission
GO:0031175: neuron projection development
R-HSA-112316: Neuronal System
GO:0120035: regulation of plasma membrane bounded cell projection organization
GO:0034330: cell junction organization
GO:0099003: vesicle-mediated transport in synapse
R-HSA-9716542: Signaling by Rho GTPases, Miro GTPases and RHOBTB3
R-HSA-9006934: Signaling by Receptor Tyrosine Kinases
GO:0007610: behavior
GO:0060627: regulation of vesicle-mediated transport
GO:0042391: regulation of membrane potential
GO:0060322: head development
R-HSA-9675108: Nervous system development
GO:0050803: regulation of synapse structure or activity
GO:0030029: actin filament-based process
GO:0098657: import into cell
R-HSA-5653656: Vesicle-mediated transport
GO:0006812: monoatomic cation transport
WP4148: Splicing factor NOVA regulated synaptic proteins
-log10 (P-value)
G
cell junction organization
actin filament-based process
Signaling by Receptor Tyrosine Kinase
regulationof cell projection organization
regulation of growth
RHO GTPase cycle
cell morphogenesis
regulation of Wnt signaling
brain development
import into cell
modulation of synaptic transmission
H
### gene overlaps SNAP-a GO categories
Synapse organization
Synaptic cell adhesion
Synaptic receptors
Synaptic vesicle
Synaptic vesicle cycle
Trans-synaptic signaling
PSD components
Neurotransmitter reuptake
Presynapse
Regulation of cholesterol biosynthesis
I
SNAP-a expression
UMAP 1
UMAP 2

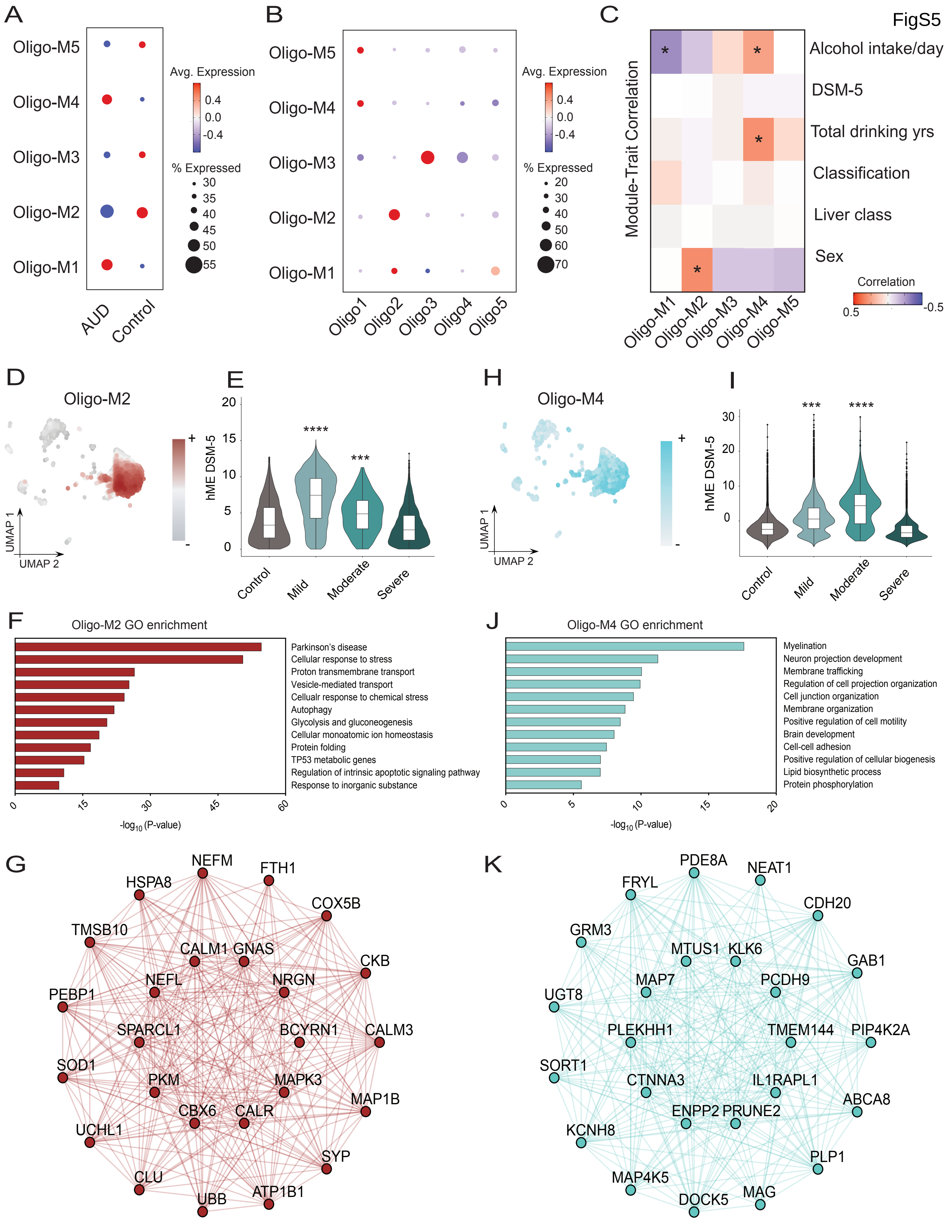
FigS5
A
Oligo-M5
Oligo-M4
Oligo-M3
Oligo-M2
Oligo-M1
AUD
Control
Avg. Expression
0.4
0.0
-0.4
% Expressed
30
35
40
45
50
55
B
Oligo1
Oligo2
Oligo3
Oligo4
Oligo5
% Expressed
20
30
40
50
60
70
C
Module-Trait Correlation
Alcohol intake/day
DSM-5
Total drinking yrs
Classification
Liver class
Sex
Correlation
0.5
-0.5
D
Oligo-M2
UMAP 1
UMAP 2
E
hME DSM-5
Control
Mild
Moderate
Severe
****
***
H
Oligo-M4
I
***
****
F
Oligo-M2 GO enrichment
Parkinson's disease
Cellular response to stress
Proton transmembrane transport
Vesicle-mediated transport
Cellualr response to chemical stress
Autophagy
Glycolysis and gluconeogenesis
Cellular monoatomic ion homeostasis
Protein folding
TP53 metabolic genes
Regulation of intrinsic apoptotic signaling pathway
Response to inorganic substance
-log10 (P-value)
J
Oligo-M4 GO enrichment
Myelination
Neuron projection development
Membrane trafficking
Regulation of cell projection organization
Cell junction organization
Membrane organization
Positive regulation of cell motility
Brain development
Cell-cell adhesion
Positive regulation of cellular biogenesis
Lipid biosynthetic process
Protein phosphorylation
G
NEFM
FTH1
HSPA8
COX5B
TMSB10
CALM1
GNAS
CKB
NEFL
NRGN
PEBP1
SPARCL1
BCYRN1
CALM3
SOD1
PKM
MAPK3
MAP1B
CBX6
CALR
UCHL1
SYP
CLU
UBB
ATP1B1
K
PDE8A
NEAT1
FRYL
CDH20
GRM3
MTUS1
KLK6
GAB1
MAP7
PCDH9
UGT8
PLEKHH1
TMEM144
PIP4K2A
SORT1
CTNNA3
IL1RAPL1
ABCA8
ENPP2
PRUNE2
KCNH8
PLP1
MAP4K5
MAG
DOCK5

FigS6

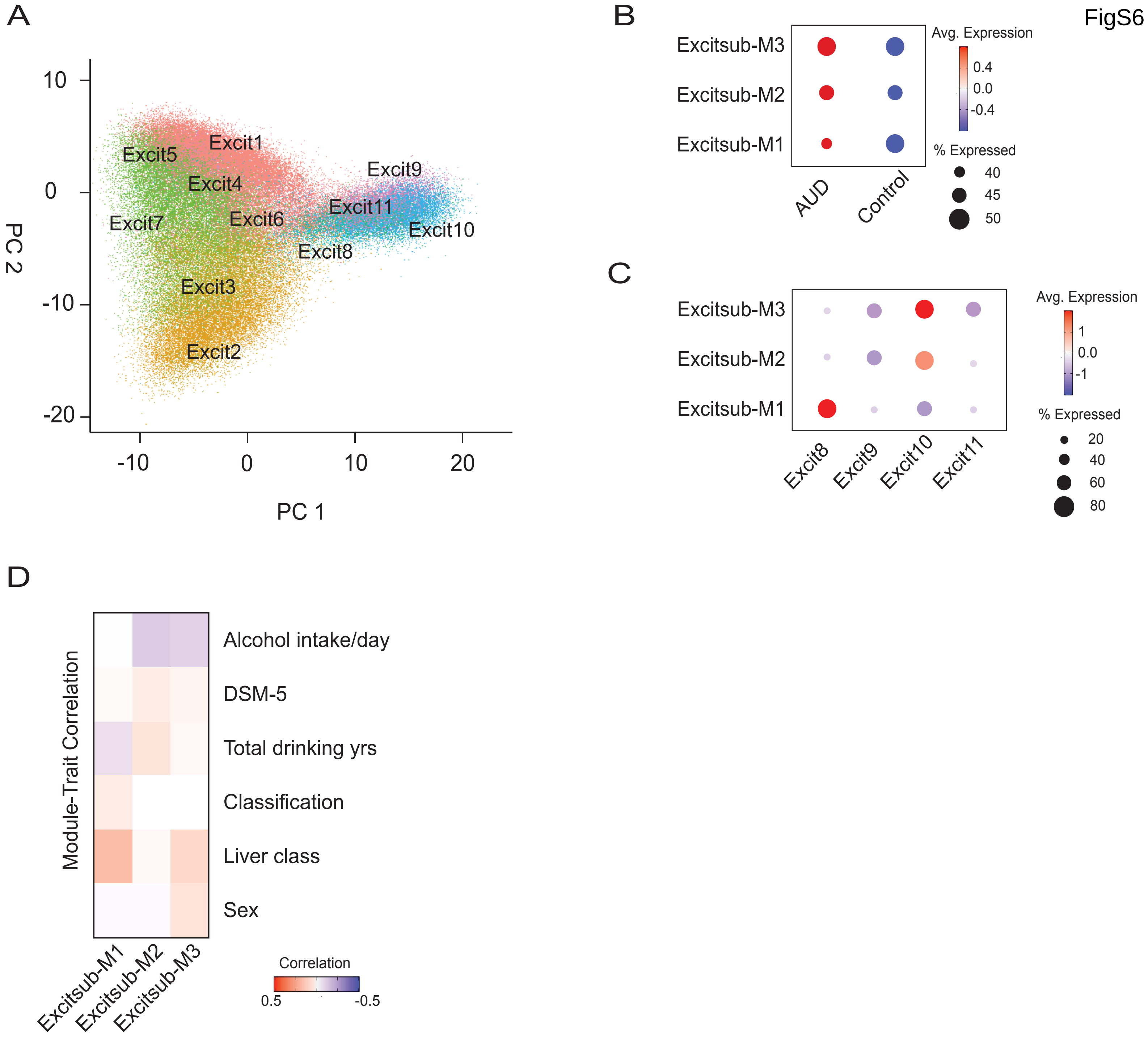
A
Excit1
Excit2
Excit3
Excit4
Excit5
Excit6
Excit7
Excit8
Excit9
Excit10
Excit11
PC 1
PC 2
10
0
-10
-20
-10
0
10
20
B
Excitsub-M3
Excitsub-M2
Excitsub-M1
AUD
Control
Avg. Expression
0.4
0.0
-0.4
% Expressed
40
45
50
C
Excitsub-M3
Excitsub-M2
Excitsub-M1
Excit8
Excit9
Excit10
Excit11
Avg. Expression
1
0.0
-1
% Expressed
20
40
60
80
D
Module-Trait Correlation
Alcohol intake/day
DSM-5
Total drinking yrs
Classification
Liver class
Sex
Excitsub-M1
Excitsub-M2
Excitsub-M3
Correlation
0.5
-0.5

Fig S7

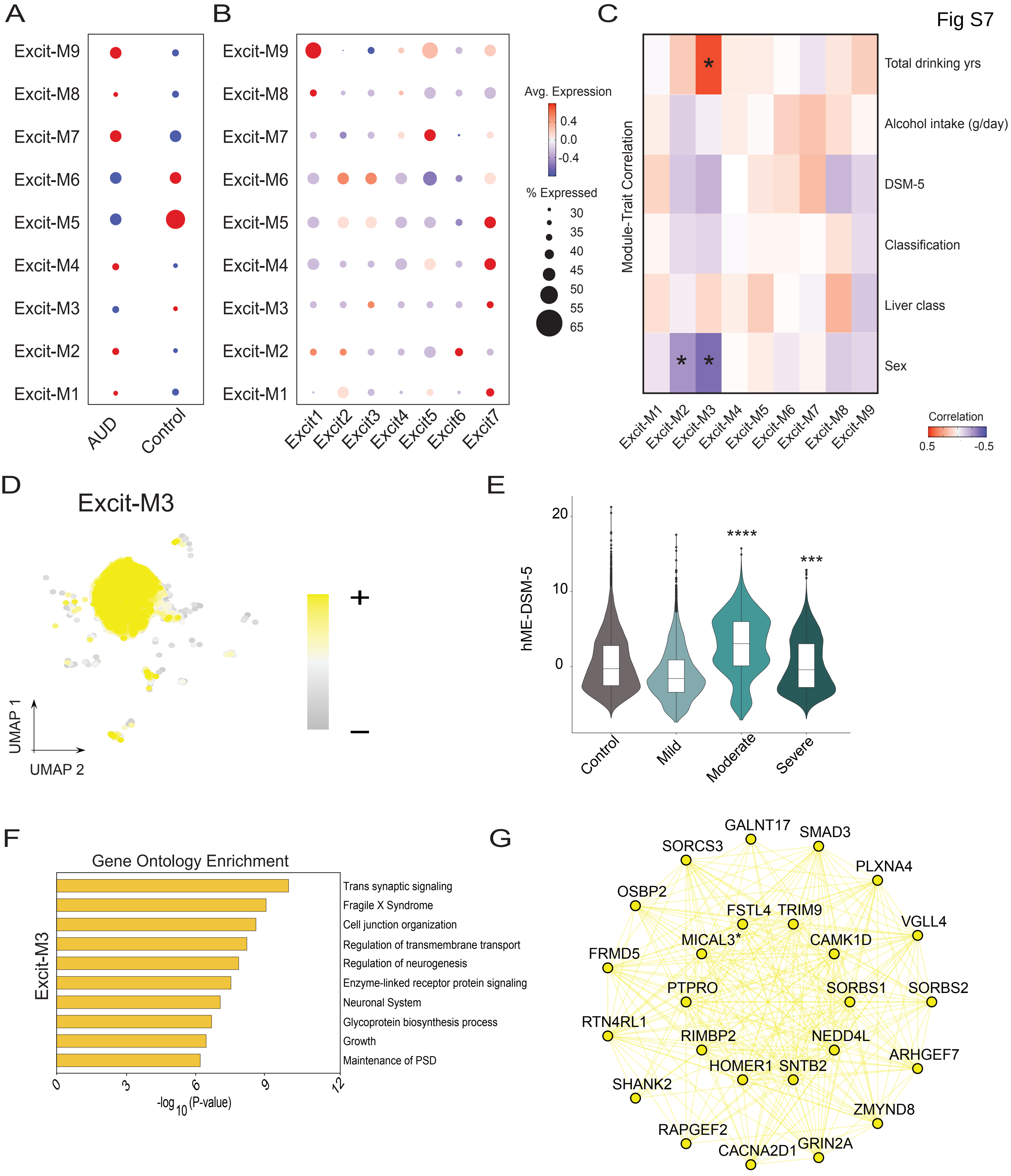
A
Excit-M9
Excit-M8
Excit-M7
Excit-M6
Excit-M5
Excit-M4
Excit-M3
Excit-M2
Excit-M1
AUD
Control
B
Excit1
Excit2
Excit3
Excit4
Excit5
Excit6
Excit7
Avg. Expression
0.4
0.0
-0.4
% Expressed
30
35
40
45
50
55
65
C
Module-Trait Correlation
Total drinking yrs
Alcohol intake (g/day)
DSM-5
Classification
Liver class
Sex
Correlation
0.5
-0.5
D
Excit-M3
UMAP 1
UMAP 2
+
–
E
hME-DSM-5
20
10
0
****
***
Control
Mild
Moderate
Severe
F
Gene Ontology Enrichment
Excit-M3
Trans synaptic signaling
Fragile X Syndrome
Cell junction organization
Regulation of transmembrane transport
Regulation of neurogenesis
Enzyme-linked receptor protein signaling
Neuronal System
Glycoprotein biosynthesis process
Growth
Maintenance of PSD
0
3
6
9
12
-log10 (P-value)
G
GALNT17
SMAD3
SORCS3
PLXNA4
OSBP2
FSTL4
TRIM9
VGLL4
MICAL3*
CAMK1D
FRMD5
PTPRO
SORBS1
SORBS2
RTN4RL1
RIMBP2
NEDD4L
ARHGEF7
HOMER1
SNTB2
SHANK2
ZMYND8
RAPGEF2
CACNA2D1
GRIN2A

Fig S8

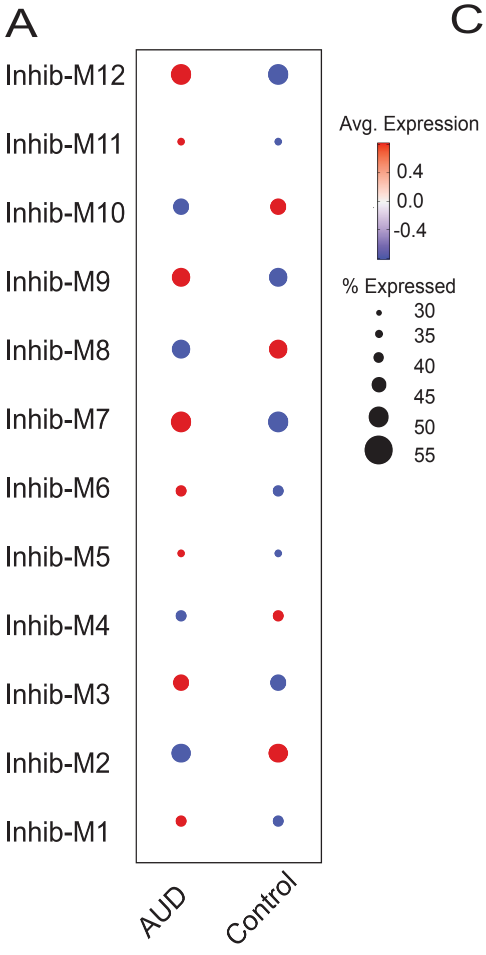

A
Inhib-M12
Inhib-M11
Inhib-M10
Inhib-M9
Inhib-M8
Inhib-M7
Inhib-M6
Inhib-M5
Inhib-M4
Inhib-M3
Inhib-M2
Inhib-M1
AUD
Control
Avg. Expression
0.4
0.0
-0.4
% Expressed
30
35
40
45
50
55


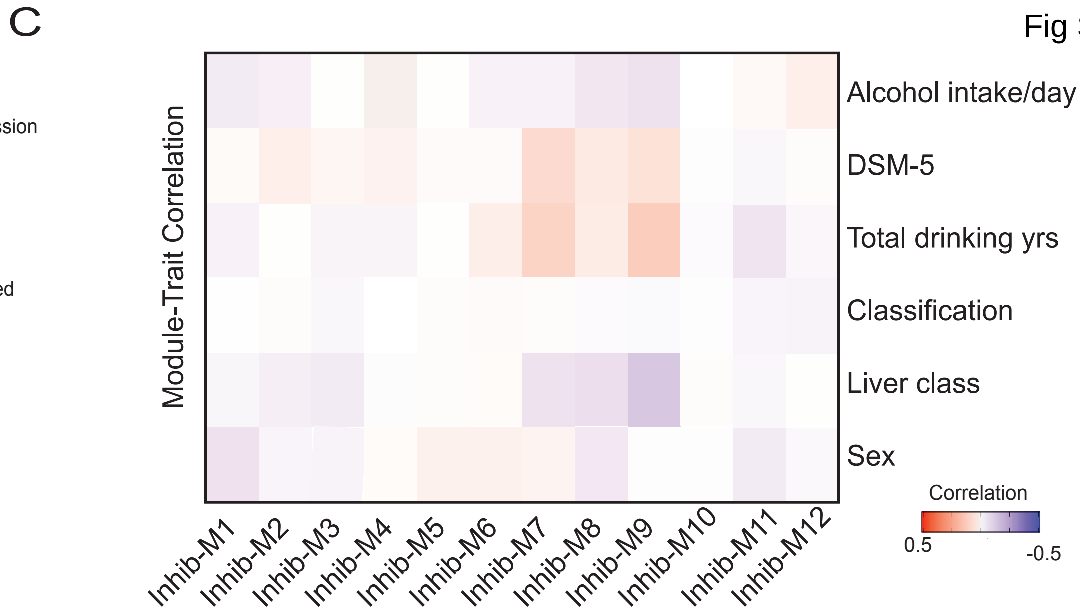

C
Module-Trait Correlation
Alcohol intake/day
DSM-5
Total drinking yrs
Classification
Liver class
Sex
Inhib-M1
Inhib-M2
Inhib-M3
Inhib-M4
Inhib-M5
Inhib-M6
Inhib-M7
Inhib-M8
Inhib-M9
Inhib-M10
Inhib-M11
Inhib-M12
Correlation
0.5
-0.5


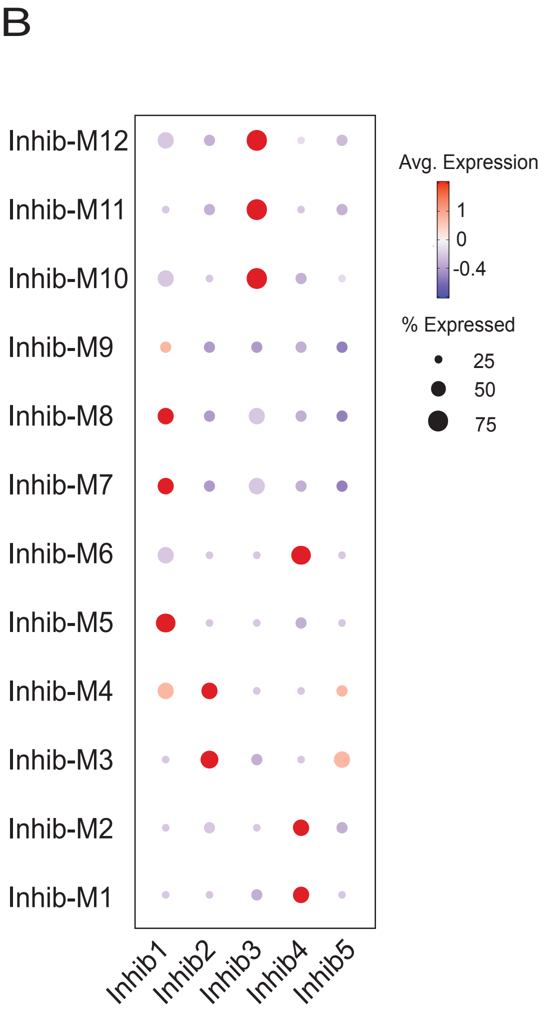

B
Inhib-M12
Inhib-M11
Inhib-M10
Inhib-M9
Inhib-M8
Inhib-M7
Inhib-M6
Inhib-M5
Inhib-M4
Inhib-M3
Inhib-M2
Inhib-M1
Inhib1
Inhib2
Inhib3
Inhib4
Inhib5
Avg. Expression
1
0
-0.4
% Expressed
25
50
75

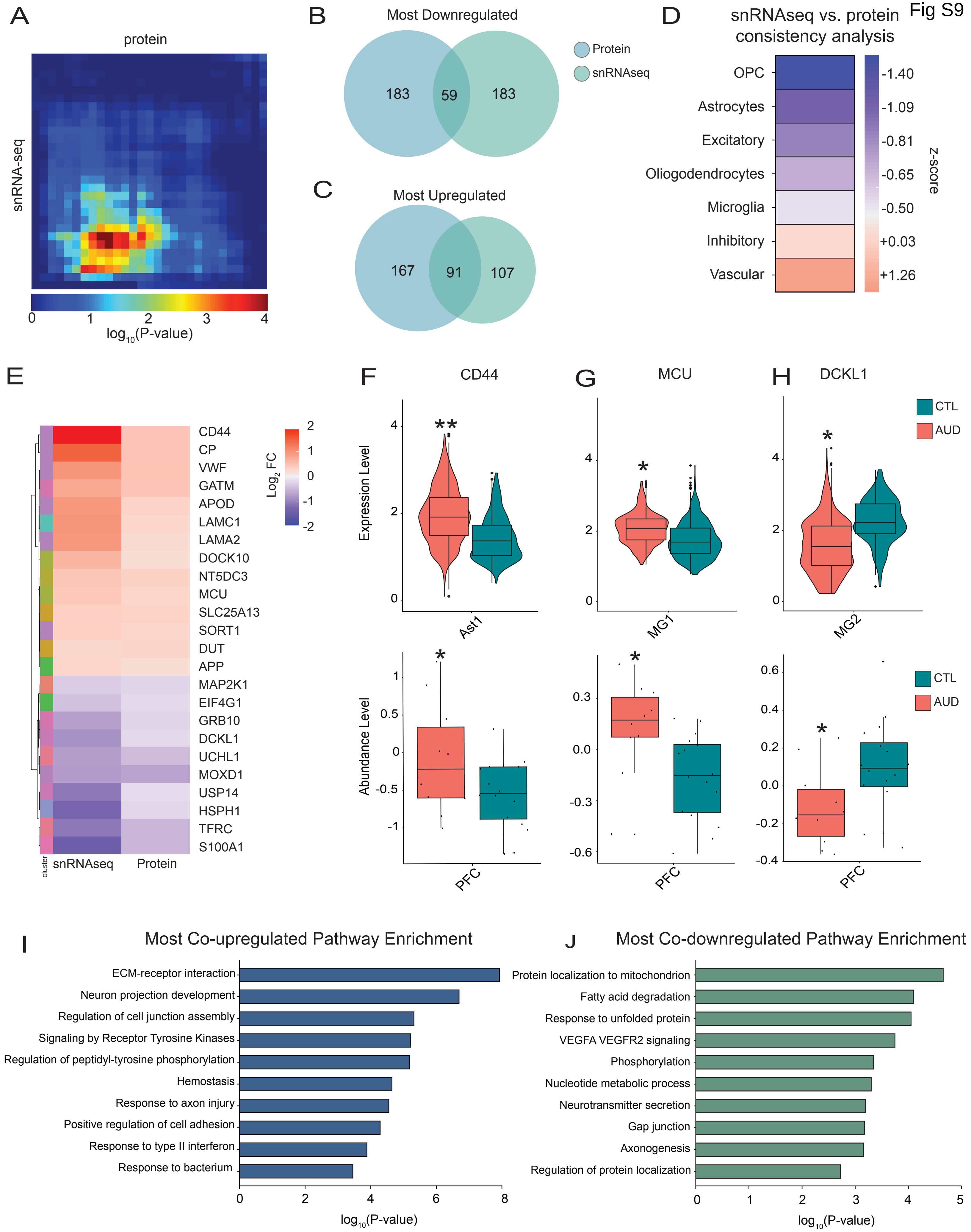
Fig S9
A
protein
snRNA-seq
0
1
2
3
4
log10(P-value)
B
Most Downregulated
183
59
183
Protein
snRNAseq
C
Most Upregulated
167
91
107
D
snRNAseq vs. protein
consistency analysis
OPC
Astrocytes
Excitatory
Oliogodendrocytes
Microglia
Inhibitory
Vascular
-1.40
-1.09
-0.81
-0.65
-0.50
+0.03
+1.26
z-score
E
CD44
CP
VWF
GATM
APOD
LAMC1
LAMA2
DOCK10
NT5DC3
MCU
SLC25A13
SORT1
DUT
APP
MAP2K1
EIF4G1
GRB10
DCKL1
UCHL1
MOXD1
USP14
HSPH1
TFRC
S100A1
cluster
snRNAseq
Protein
Log2 FC
F
CD44
G
MCU
H
DCKL1
CTL
AUD
Expression Level
Ast1
MG1
MG2
Abundance Level
PFC
I
Most Co-upregulated Pathway Enrichment
ECM-receptor interaction
Neuron projection development
Regulation of cell junction assembly
Signaling by Receptor Tyrosine Kinases
Regulation of peptidyl-tyrosine phosphorylation
Hemostasis
Response to axon injury
Positive regulation of cell adhesion
Response to type II interferon
Response to bacterium
log10(P-value)
J
Most Co-downregulated Pathway Enrichment
Protein localization to mitochondrion
Fatty acid degradation
Response to unfolded protein
VEGFA VEGFR2 signaling
Phosphorylation
Nucleotide metabolic process
Neurotransmitter secretion
Gap junction
Axonogenesis
Regulation of protein localization
log10(P-value)
